# Restriction on Ku’s Inward Translocation Caps Telomere Ends

**DOI:** 10.1101/2024.12.22.629684

**Authors:** Stefano Mattarocci, Sonia Baconnais, Florian Roisné-Hamelin, Sabrina Pobiega, Olivier Alibert, Vincent Morin, Alice Deshayes, Xavier Veaute, Virginie Ropars, Maelenn Chevreuil, Johannes Mehringer, Didier Busso, Gerard Mazon, Paloma Fernandez Varela, Éric Le Cam, Jean-Baptiste Charbonnier, Philippe Cuniasse, Stéphane Marcand

## Abstract

Safeguarding chromosome ends against fusions via nonhomologous end joining (NHEJ) is essential for preserving genome integrity. Paradoxically, the conserved NHEJ core factor Ku binds telomere ends. How it is prevented from promoting NHEJ remains unclear, as does the mechanism that allows Ku to coexist with telomere-protective DNA binding proteins, e.g., Rap1 in *Saccharomyces cerevisiae*. Here, we reveal a direct role for Rap1 in the inhibition of Ku’s NHEJ function at telomeres. A single Rap1 molecule bound near a DNA end inhibits NHEJ *in vivo* without disrupting Ku presence. Consistent with this, Rap1 and Ku form a complex on short DNA duplexes *in vitro*. Cryo-EM and molecular modelling analysis of this complex shows that Rap1 obstructs Ku’s inward translocation on DNA – an essential step for NHEJ at broken ends. Nanopore sequencing of telomere fusions confirms the importance of this pathway in protecting native telomere ends. Collectively, our findings uncover a mechanism of telomere end protection mediated by restricting Ku’s inward translocation, a functional switch that prevents promiscuous NHEJ repair at telomeres.

## INTRODUCTION

Telomeres maintain genome stability by protecting native chromosome ends from the DNA damage checkpoint and repair pathways that act on broken ends ^1 2^. In most eukaryotes, telomeric DNA consists of tandemly repeated sequences bound by high-affinity telomere proteins. This repetitive protein occupancy is a distinctive feature of telomeres and is critical for their functions. Short telomeres, with few DNA-bound telomere proteins, are more susceptible to fusions, checkpoint activation, and telomere loss ^3 4 5^. They are also more likely to be elongated by telomerase, which restores full protection ^6 7^. This strong length-dependence suggests that, within telomeres, end-distal and end-proximal telomere proteins jointly contribute to telomere functions. However, it is unclear whether their contributions are equivalent and/or if end-proximal telomere proteins play specific roles due to their position at the forefront to encounter the DNA damage checkpoint and repair proteins that bind to DNA ends ^8 9^.

The NHEJ Ku70-Ku80 heterodimer (Yku70-Yku80 in *S. cerevisiae*, and referred to hereafter as Ku) is the first factor to bind broken ends in all eukaryotes ^10^. Ku has a ring-shaped structure that encircles DNA double-stranded ends with high-affinity, but without sequence specificity ^10 11 12 13 14 15^. Ku protects broken ends from resection initiation and recruits the NHEJ-specific ligase 4 along with its associated protein partners. Together, they mediate end synapsis followed by end ligation, a process that requires Ku to translocate inward by approximately 25 base pairs on the DNA, thereby making the ends accessible to the ligase ^10 15 16 17 18^. Paradoxically, Ku constitutively binds telomere ends and protects them from uncontrolled resection without engaging NHEJ, a feature conserved across yeast, mammals and plants ^19 20 21 22 23 24 25 26 27 28 29 30^. How Ku’s roles in promoting NHEJ are so effectively suppressed at telomeres remains unknown. It is also unclear whether Ku and end-proximal telomere proteins compete or cooperate for DNA binding at telomere ends.

We addressed these issues in *S. cerevisiae*. In this species, the protein Rap1 directly binds telomere repeats with high-affinity and protects telomeres against NHEJ-dependent fusions and uncontrolled end resection through multiple mechanisms ^4 24 31 32 33 34 35 36 37 38^ (**Fig. S1-2**). The reliance on multiple pathways acting in synergy is a conserved feature of telomere end protection ^23 30 39 40 41 42 43^. Each can provide significant, non-redundant and potentially similar levels of protection ^4 37 38^ (**Fig. S1**). Understanding the molecular bases of telomere stability requires identifying and deciphering each of these pathways. Here, we uncover a new mechanism of telomere end protection that relies specifically on the end-proximal, telomere-bound Rap1 protein. This mechanism operates independently of telomere length and acts by restricting Ku’s inward translocation, thereby preventing promiscuous NHEJ repair at telomeres.

## RESULTS

### NHEJ inhibition at a broken end by a single Rap1

To assess the potential contribution of the end-proximal telomere-bound Rap1 protein to telomere protection, we investigated whether a single Rap1 molecule when bound close to a broken end, could significantly antagonise NHEJ repair. Within *S. cerevisiae* telomere sequences, the motif ^5’^GGTGTGTGGGTGTG^3’^ matches the Rap1 consensus binding site ^32 44^ (**Fig. 1A**, **Fig. S1A**). Rap1 binds this sequence with subnanomolar affinity *in vitro* ^34 45^ (**Fig. S3**). We therefore tested whether a single copy of this Rap1 site could inhibit NHEJ in an assay where survival following continuous expression of a site-specific endonuclease serves as a proxy for NHEJ repair efficiency ^33 37^ (**Fig. 1B**).

**Figure 1.**
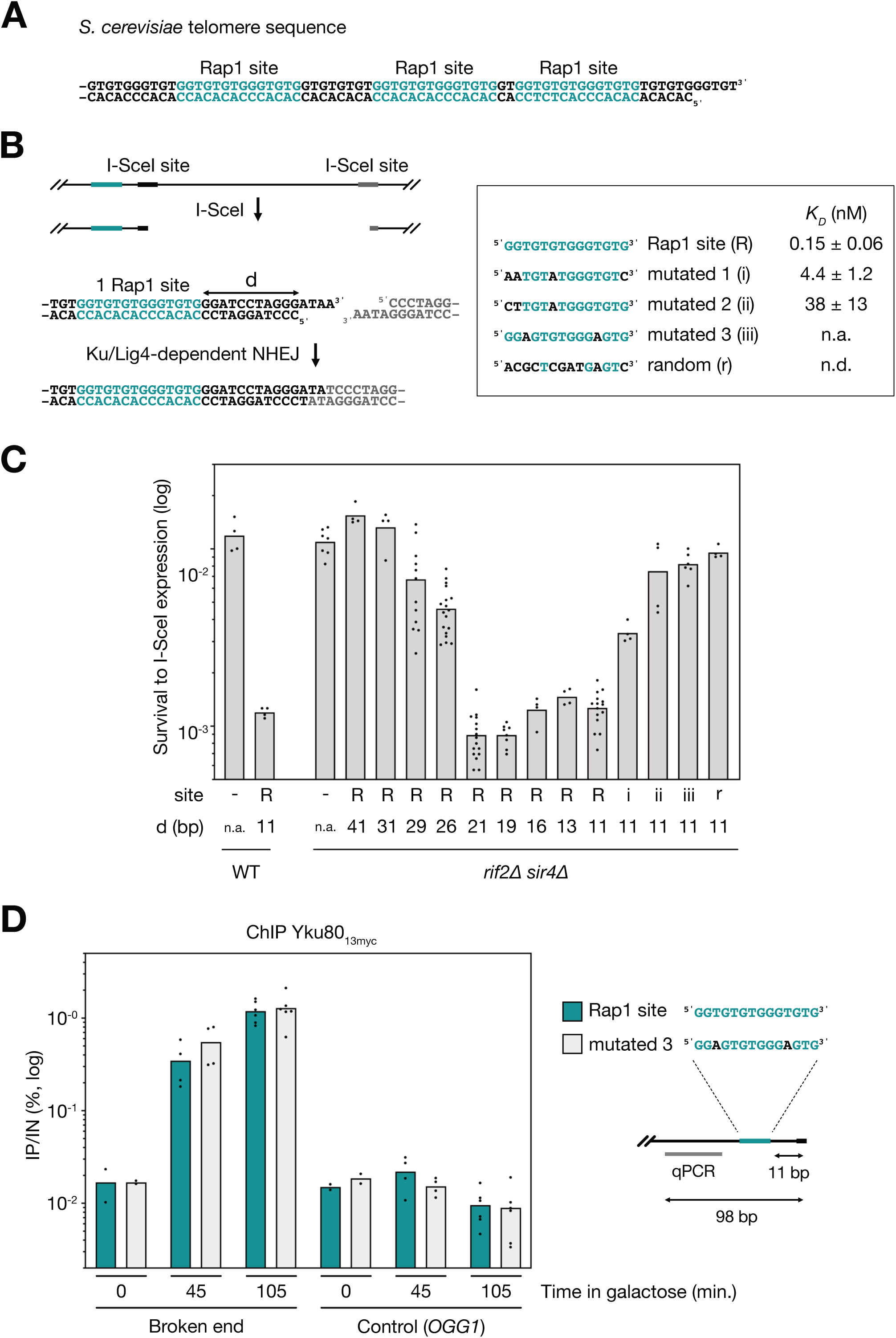
Inhibition of NHEJ by a single Rap1 site at broken ends. (**A**) Example of an *S. cerevisiae* telomere sequence with the Rap1 sites highlighted in teal. (**B**) Diagram of the I-SceI assay used to estimate NHEJ efficiency. Two inverted I-SceI sites were inserted at the endogenous *URA3* gene. Survivors to continuous I-SceI expression have eliminated the I-SceI sites by fusing the distal broken ends ^33 37^. A single Rap1 site was inserted next to one of the I-SceI sites. Inset: sequences of the tested mutated sites, with their respective affinities for Rap1 *in vitro* (*K_D_* values from **Fig. S3**). (**C**) NHEJ inhibition by a single Rap1 targeted at a broken end in wild-type (WT) cells and cells lacking Rif2 and Sir4 (*rif2Δ sir4Δ*). Mean from independent cell cultures. Statistical analysis are shown in **Table S1**. *(d)* number of base pairs separating the edge of the Rap1 site from the broken end, *(-)* no insert, *(R)* 1 Rap1 site, *(i to iii)* mutated sites 1 to 3, *(r)* random 14 bp sequence. (**D**) Rap1 binding does not prevent Ku recruitment at a broken end. Yku80 presence at a broken end (after 45 and 105 min of I-SceI induction by galactose) with a Rap1 site (teal) or a mutated site (light gray) was determined by ChIP (time 0: uncut control). Control qPCR at the *OGG1* locus. Rap1 site was positioned 11 bp from the broken end. Cells were arrested in G1 to limit end resection and therefore preserve Rap1 and Ku binding. Quantification of immunoprecipitated DNA (IP) relative to the input DNA (IN) is shown. Means from independent samples. The control was performed within the *OGG1* coding sequence.

The insertion of a Rap1 site positioned 11 bp from the double-stranded end of the broken end led to significant inhibition of NHEJ repair (d = 11 bp, **Fig 1C**). The strength of this inhibition was comparable in the presence or absence of Rif2 and Sir4, two Rap1 co-factors previously shown to inhibit NHEJ at telomeres ^4 33 36 38 37^. This rules out any contribution of Rif2 and Sir4 in the NHEJ inhibition mediated by an isolated Rap1 molecule and supports the hypothesis that a previously unidentified mechanism must be at play. A Rap1 site located more than 30 bp from the end had no impact on NHEJ repair, indicating that the action of Rap1 on NHEJ is short-range (**Fig. 1C**, statistical analysis in **Table S1**). Furthermore, mutations in the Rap1 site that lower the affinity for Rap1 *in vitro* ^34^ (**Fig. 1B**, **Fig. S3**) weakened NHEJ inhibition (**Fig. 1C**). Interestingly, native Rap1 sites that diverge from the consensus Rap1 telomere site were also less effective at blocking NHEJ when inserted at a broken end (**Fig. S4-S5**). Together, these results show that a single Rap1 molecule, tightly bound to DNA ends, can significantly protect against fusion by NHEJ.

### Rap1 and Ku co-binding at DNA ends

Can a single Rap1 block NHEJ at a broken end by preventing Ku binding? Using a ChIP approach, we found that Rap1 binding 11 bp from a broken end did not inhibit Ku’s recruitment (**Fig. 1D**, **Fig. S6**), which is consistent with the co-presence of Rap1 and Ku at telomeres *in vivo*. We next explored the conditions under which Rap1 and Ku could simultaneously bind the same DNA end *in vitro*. Binding experiments using purified proteins and Electrophoretic Mobility Shift Assays (EMSA) show that, individually, Rap1 and Ku bound specifically to a 17-bp DNA duplex containing a single Rap1 site and only a 1 bp extension downstream of the site (**Fig. 2A**, **Fig. S7A-B**). However, when incubated together with the same DNA duplex, their binding appeared mutually exclusive (**Fig. 2A**, **Fig. S7C**), suggesting that Rap1 binding on this DNA sterically hinders Ku’s ability to bind to the DNA end.

**Figure 2.**
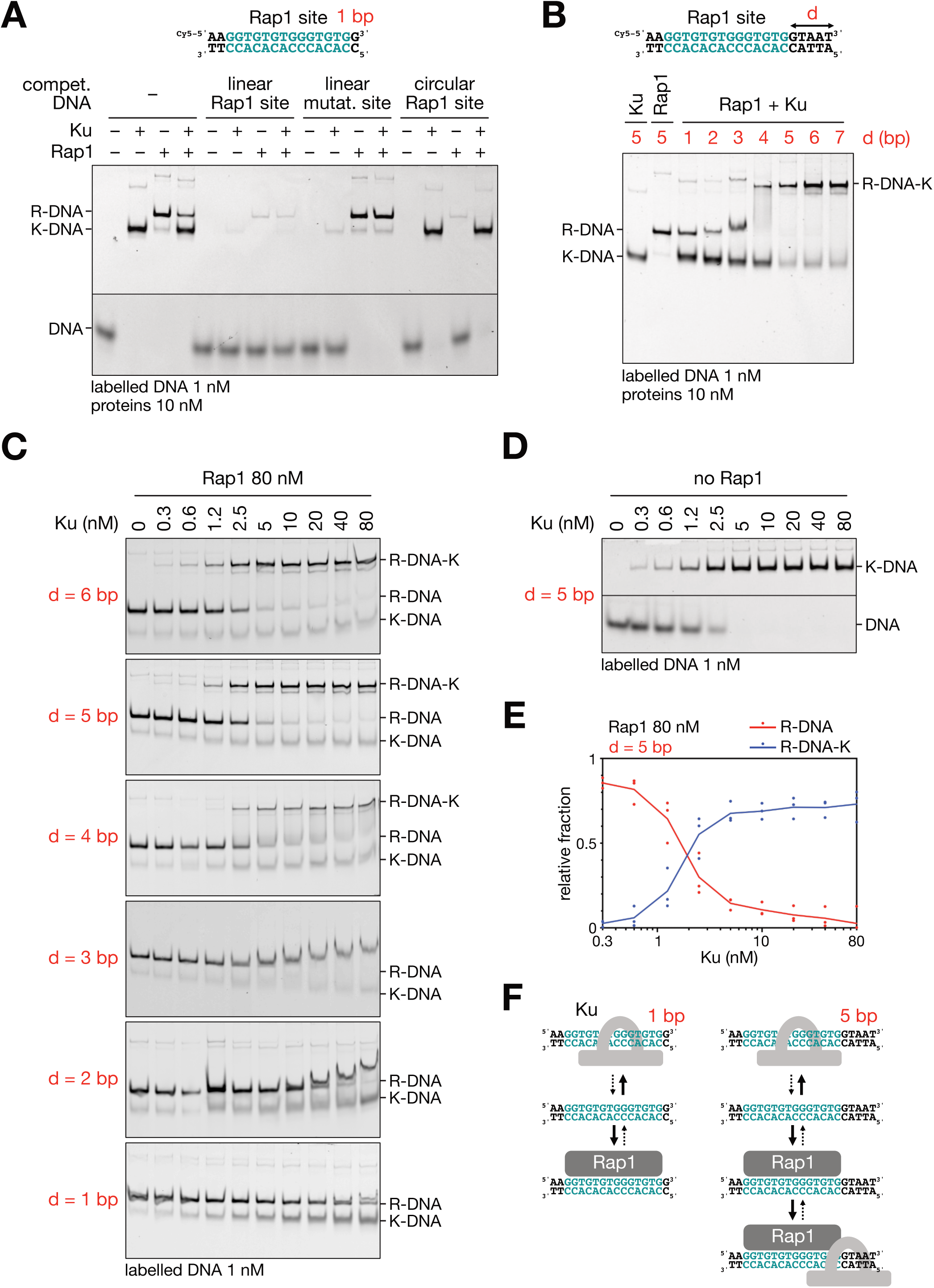
Ku binding on Rap1-DNA complexes. (**A**) Binding of Ku and Rap1 to a short DNA duplex containing a Rap1 site and a 1 bp downstream extension (1 nM). Unlabelled competitor DNA: (i) short linear duplex with a Rap1 site and a 1 bp downstream extension (titrates Rap1 and Ku, 200 nM), (ii) short linear duplex with a mutated site (mutated 3 in Fig. 1 and Fig. S3) and a 1 bp downstream extension (titrates Ku only, 200 nM), circular plasmid with 16 Rap1 sites in tandem ^45^ (titrates Rap1 only, 20 nM). *(R)* represents Rap1, *(K)* represents Ku. (**B**) Binding of Ku and Rap1 to DNA duplexes with downstream extensions ranging from 1 to 7 bp (1 nM). *(d)* refers number of base pairs separating the edge of the Rap1 site from the duplex end. (**C**) Representative titration of Ku binding to DNA duplexes with downstream extensions of 1 to 6 bp (1 nM) in the presence of excess Rap1 at a fixed concentration of 80 nM. (**D**) Representative titration of Ku binding to DNA in the absence of Rap1. (**E**) Quantified EMSA data showing Ku association on Rap1-bound DNA with a downstream extension of 5 bp (1 nM), as in (C). Means from 3 independent experiments. (**F**) Interpretative schematic illustrating the mutual exclusion or co-binding of Rap1 and Ku on short duplex DNA with a 1 or 5 bp extension downstream of the Rap1 site.

We then investigated the length of DNA downstream of the Rap1 binding site that would be sufficient to support Ku binding to Rap1-bound DNA duplexes. DNA extensions of 4 bp or more shifted the equilibrium towards the formation of larger complexes, accommodating both Rap1 and Ku on the same DNA molecule (d ≥ 4 bp, **Fig. 2B**, **Fig. S7D**). This suggests that these few base pairs downstream of Rap1 are sufficient to enable Ku binding to Rap1-DNA complexes, forming ternary complexes. When we progressively increased the concentration of both Rap1 and Ku (up to 80 nM each), these ternary complexes became predominant (**Fig. S8**). To better estimate the binding of Ku on the Rap1-bound DNA ends, we fixed Rap1 concentration at 80nM and increased progressively Ku concentration (**Fig. 2C**, to compare to Ku binding on free DNA shown in **Fig. 2D**). We observed that the association between Ku and Rap1-DNA complexes with a 5 bp extension predominates at Ku concentrations above 2 nM (**Fig. 2C&E**), suggesting an affinity in the low nanomolar range or less, close to the affinity of Ku for free DNA ends (**Fig. 2D**, **Fig. S3**). On Rap1-bound DNA with shorter extensions of 3 bp or less, we note that Ku may still transiently bind at higher protein concentrations but dissociates during migration in the gel (**Fig. 2B-C**). Thus, the DNA length downstream of the Rap1 binding site is a key factor in determining Ku association on Rap1-DNA complexes. To corroborate these findings, we employed Mass Photometry. This method confirmed that Rap1 antagonises Ku binding on DNA duplexes with 2 bp downstream of the Rap1 site, but not on those with 6 bp, which is sufficient to allow Ku binding on Rap1-DNA complexes (**Fig. S9**).

The DNA binding domain of Rap1 (Rap1_DBD_) comprises two Myb-like Helix-Turn-Helix subdomains (Myb_1_ and Myb_2_), each engaging with a 5-6 bp hemi-site ^32^. Mutations in either hemi-site impaired both Rap1’s interaction with the DNA duplex and the formation of Rap1-DNA-Ku complexes (**Fig. S10**), indicating that Rap1 engages with both hemi-sites in these complexes and ruling out the possibility of a partial Rap1 interaction that might leave additional free DNA for Ku binding ^46^. 3’ single-stranded overhangs of up to 12 nucleotides did not prevent Ku from binding to Rap1-DNA complexes (**Fig. S11)**, suggesting that Ku is not obstructed by the short 3’ overhangs found at native yeast telomeres ^31^. Collectively, these findings show that Ku associates with Rap1-bound DNA ends when there is a double-stranded DNA extension of 4 bp or more downstream of Rap1. With shorter DNA extensions, Rap1 and Ku binding tend to become mutually exclusive (**Fig. 2F**).

### Structural analysis of Rap1 ability to accommodate Ku at DNA ends

The apparent threshold length for Ku binding to Rap1-bound DNA ends (4 bp, **Fig. 2**) is unexpectedly short compared to the minimal number of base pairs typically bound by Ku, as seen in resolved structures (>10 bp) ^10 11 13 14^. To address this apparent paradox, we utilised molecular modelling and cryo-electron microscopy. Initially, we built a model of the ternary Rap1-DNA-Ku complex using the crystal structures of *S. cerevisiae* Ku ^13^ and of *S. cerevisiae* Rap1_DBD_ bound to a DNA duplex ^47^. In this model, the 21 bp DNA duplex includes a 5 bp extension downstream of the 14 bp Rap1 site. To perform cryo-EM, we incubated the same 21 bp DNA duplex with Rap1 followed by Ku and then purified and vitrified the complexes. The collected data allowed us to obtain a map of the ternary Rap1-DNA-Ku complex at an overall resolution of 3.1 Å. The initial model was further refined through molecular dynamics flexible fitting constrained by the cryo-EM map (**Fig. 3A-B**, **Fig. S12-S13**, **S14A**, **S15**, **S16A**, **Table S2**, **Movie S1**).

**Figure 3.**
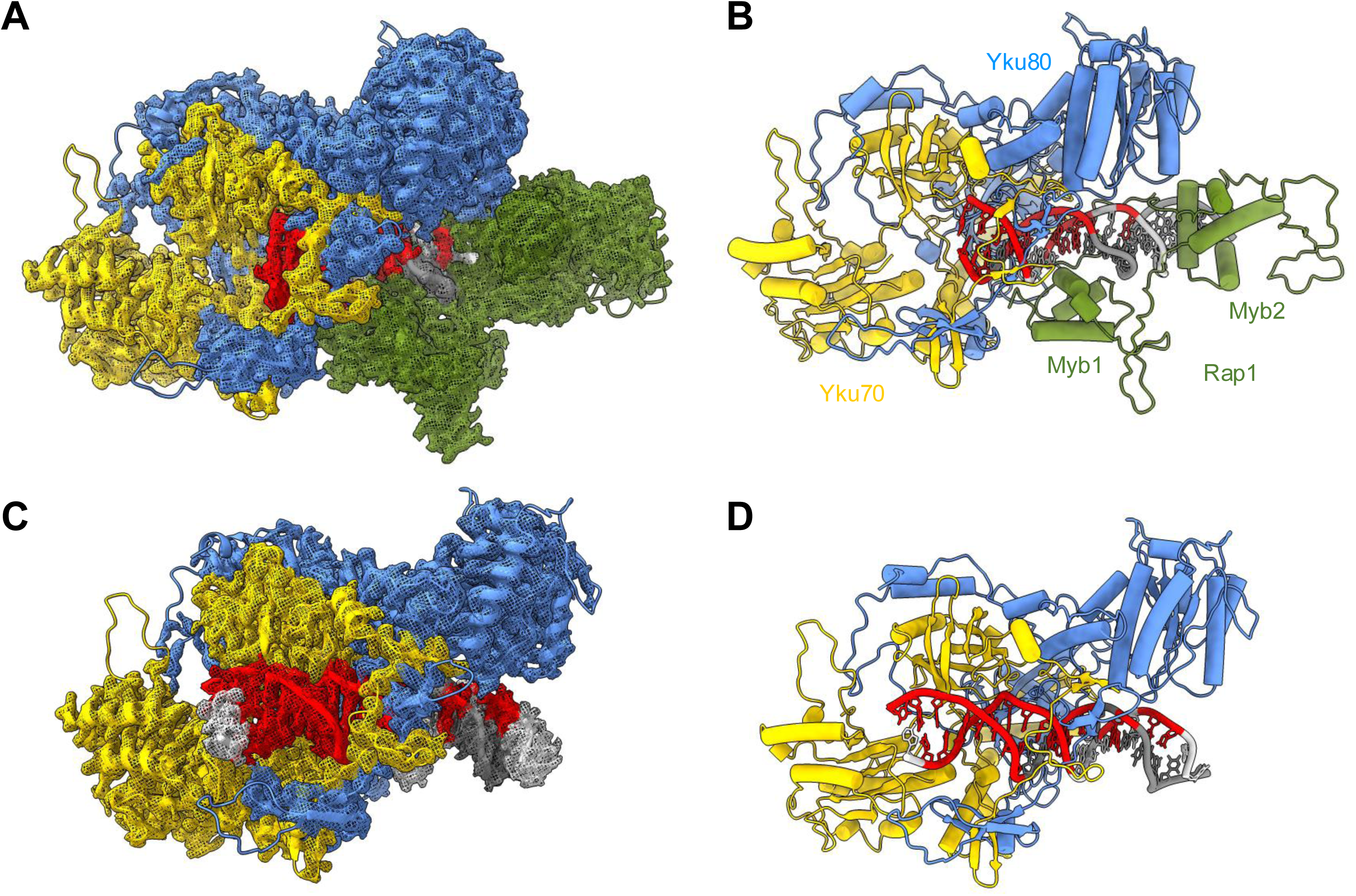
Structural analysis of Rap1’s ability to restrict Ku’s inward translocation on DNA. Structure of the Rap1_DBD_-DNA-Ku ternary complex (**A-B**) and the DNA-Ku binary complex (**C-D**) shown in cartoon representation. Yku70 is shown in gold, Yku80 in cornflower blue, Rap1_DBD_ in green, G-rich DNA strand in light grey and C-rich DNA strand in grey, with nucleotides located within 4.0 Å of Ku marked in red (both strands). The cryo-EM maps are shown in mesh (**A** and **C**) and coloured according to the atom proximity in the refined model. At the thresholds used, electron density is not visible in regions corresponding to residues 194-211 of Yku70, 165-171, 287-301, 462-466 of Yku80 and residues 429-433, 480-506 and 598-601 of Rap1_DBD_ in the Rap1_DBD_-DNA-Ku complex (**A**) and regions corresponding to 200-211 of Yku70, 95-104, 165-171, 263-270, 286-300 and 579-587 of Yku80 in the DNA-Ku complex (**C**).

The resolved portion of Rap1 is restricted to its DNA binding domain, which aligns closely with the Rap1_DBD_-DNA crystal structure ^47^ (**Table S2**). Rap1 engages the DNA major groove via its two Myb-like subdomains, along with a loop at the C-terminal region of Myb_2_ (residues 565 to 601), which wraps around the DNA duplex and contacts the Myb1 subdomain, effectively clamping the molecule ^32 47^ (**Fig. S15**). The Ku structure in the ternary complex also closely resembles the *S. cerevisiae* Ku crystal structure ^13^ (**Table S2**, **Fig. S17A**). Ku’s interaction with DNA involves five base pairs at the DNA extremity (the extension downstream of the Rap1 site), which are engaged into the Ku beta-bridge (handle), in addition to eight more base pairs that also interact with Rap1 Myb_1_ subdomain (**Fig. 3A-B**, Ku-interacting nucleotides in red). This is permitted because the two proteins bind different sides of the same DNA double helix (**Fig. S18**) and display a remarkable shape compatibility (**Fig. 3A-B**, **S15**). Beyond the beta-bridge, Ku does not contact the DNA (**Fig 3A-B**, **S15**). In total, Ku engagement with 13 bp of DNA is consistent with previous estimates for Ku binding to duplex DNA ^10 11 13 14^.

Although Rap1 and Ku bind very closely to each other, contacts between Rap1 and Ku (distance < 4 Å) are restricted to two small regions (**Fig. S19**). One involves seven residues of the Rap1 Myb_2_ subdomain and a six-residue loop in Yku80 (patch 1), while the other involves four residues from in the Rap1 C-terminal wrapping loop and five residues at the edge of Yku80 and Yku70 (patch 2) (**Fig. S19**). Of these residues, only four are compatible with hydrogen bond or salt bridge distances (a salt bridge between Yku80 R124 and Rap1 E572, and a hydrogen bond between Yku80 Q122 and Rap1 Y552, **Fig. S19**). Two of these residues are poorly conserved across yeast species (Rap1 E572 and Yku80 Q122, **Fig. S20A**), suggesting that specific interaction between the two proteins plays a minimal role in the ternary complex formation. Supporting this hypothesis, mutations in the Yku80 residues that contact Rap1 (including Q122 and R124) did not impair Rap1’s ability to inhibit NHEJ at a broken end (**Fig. S20B**).

Among the particles used to generate the cryo-EM map of the ternary complex, we also identified particles corresponding to a DNA-Ku binary complex. The map of this complex was determined at an overall resolution of 2.9 Å and was used to refine an initial DNA-Ku model by molecular dynamics flexible fitting and real-space refinement (**Fig. 3C-D**, **Fig. S12-S13**, **S14B**, **S16B**, **Table S2**, **Movie S2**). As observed in the ternary complex, the Ku structure in the DNA-Ku complex closely resembles the *S. cerevisiae* Ku crystal structure ^13^ (**Table S2**, **Fig. S17B**). In the binary complex, the 21 bp DNA duplex is well engaged on both sides of the Ku handle. The interface between Ku and DNA is more extensive than in the ternary complex, involving 18 bp rather than 13 bp (**Fig. 3C-D**, Ku-interacting nucleotides in red, **Table S2**). The distinct positioning of the DNA duplex in Ku between the binary and ternary complexes show that, in the absence of Rap1, the DNA duplex more fully occupies the DNA-binding site of Ku. This indicates that Rap1 limits how far DNA can engage into the Ku ring. In other words, Rap1 binding to DNA restricts by steric hindrance Ku’s inward translocation on the DNA following Ku’s initial binding at a DNA end. Since adequate Ku’s inward translocation is essential for its role in NHEJ ^15 18^, Rap1’s inhibition of this process may safeguard telomeres from NHEJ.

### Significance at telomeres of Rap1 ability to restrict Ku’s inward translocation

If Rap1’s control of Ku’s inward translocation is important for telomere protection, weakening this control should result in an increase in telomere fusions. The absence of a significant specific interaction between Rap1 and Ku prevents the design of specific mutants to investigate the function of the ternary complex using conventional genetics (**Fig. S19, S20**). To overcome this limitation, we developed an alternative approach based on our observation that Ku’s binding to Rap1-bound DNA ends is influenced by the length of DNA downstream of Rap1.

When Rap1 is positioned at an optimal distance from the DNA end, Ku can bind these ends while Rap1 effectively blocks its inward translocation, thereby protecting these telomeres. However, when Rap1 is positioned too close to the DNA end, the binding of Rap1 and Ku tends to become mutually exclusive, as observed *in vitro* on short DNA duplexes (**Fig. 2**, **Fig. S7**). In such cases, Ku may bind in place of Rap1 and subsequently translocate inward to a position that supports NHEJ. If this hypothesis is correct, telomeres where the most-end proximal Rap1 is located very close to the end should be more prone to fusion, all else being equal. By sequencing telomere fusions and determining the position of Rap1 sites relative to the fusion points, we can test this prediction and, in turn, the significance of Rap1’s ability to regulate Ku.

For this approach, telomere fusions were PCR-amplified from Rap1-proficient cells lacking Rif2 and Sir4 and sequenced using Nanopore sequencing (**Fig. 4A**, **Fig. S21A-B**). We used cells lacking Rif2 and Sir4 for two reasons: (1) telomere fusions in wild-type cells are too rare to be amplified ^4 33 36^, and (2) the absence of Rif2 and Sir4 maximises reliance on the remaining pathways established by Rap1, which supports our aim of correlating Rap1 positioning with its efficiency in protecting telomeres. We mapped the telomere sequences, their fusion points, and the position of the Rap1 sites relative to these points. We then calculated the frequency of Rap1 sites (i.e., their relative occurrence) as a function of their distance from the fusion points.

**Figure 4.**
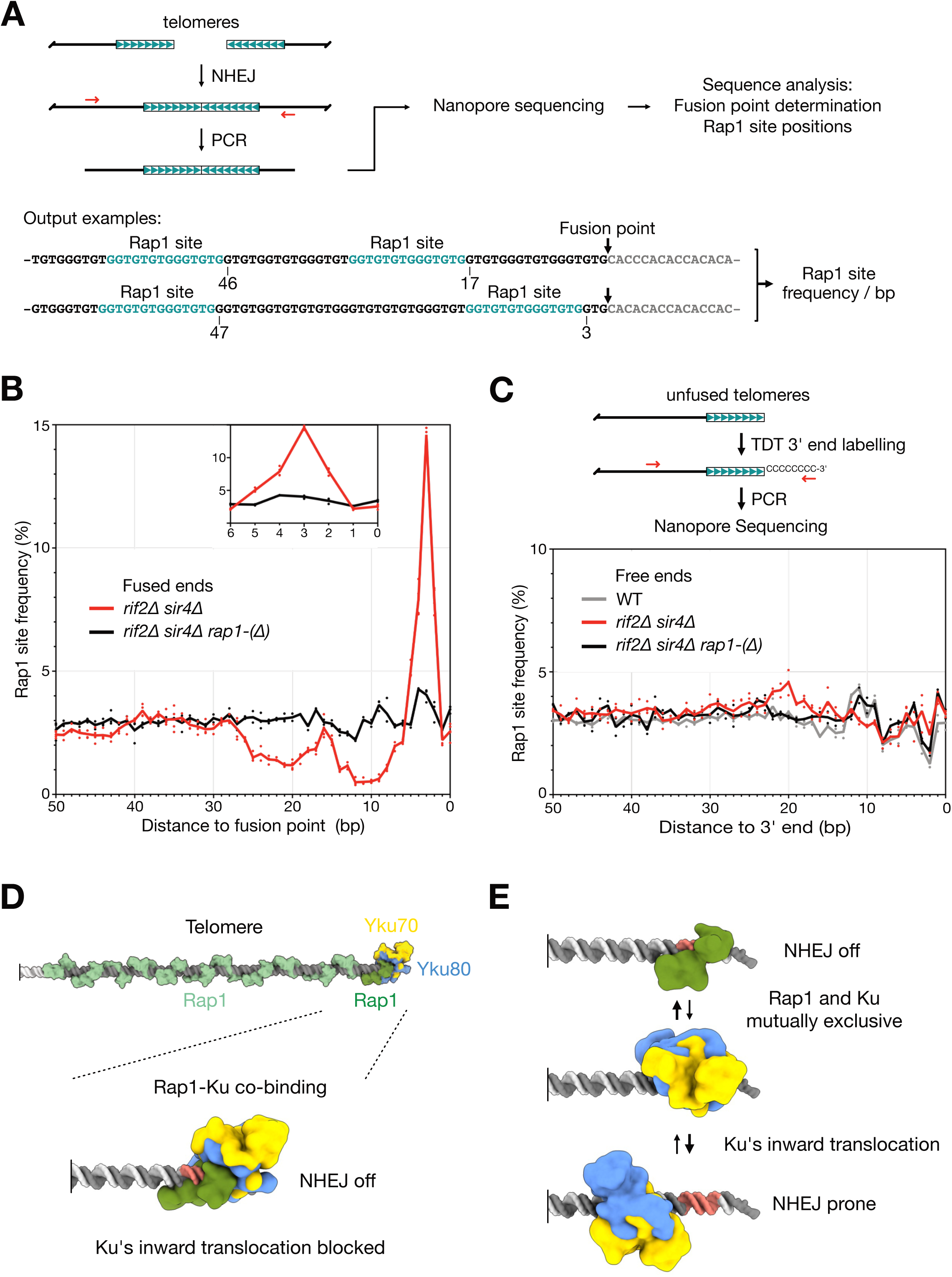
End-proximal Rap1 site position impacts telomere end protection. (**A**) Schematic of the approach used to determine Rap1 site positions at fused telomeres. Two examples of sequenced telomere-telomere fusions are shown. Rap1 sites are highlighted in teal. (**B**) Rap1 site frequency at fused telomere ends. G-rich strand sequences. Fusion points at position 0. Means from independent cell cultures and sequencing experiments. (**C**) *Top* - schematic of the approach used to determine Rap1 site positions at unfused telomeres. X and Y’ telomere 3’ ends are labelled by TDT prior to PCR amplification ^52^. *Bottom* - Rap1 site frequency at unfused telomere ends. G-rich strand sequences. Telomere 3’ ends at position 0. (**D**) Model for telomere capping by the end-proximal Rap1. Yku70 is shown in gold, Yku80 in cornflower blue, end-proximal telomere-bound Rap1_DBD_ in green, other telomere-bound Rap1_DBD_ in light green, telomere DNA in grey, subtelomere DNA in light grey, with the Rap1 site highlighted in salmon (both strands). For simplicity, other telomere components, such as NHEJ inhibitors Rif2 and Sir4, are not represented. Ku binding and the restriction of its inward translocation by Rap1 predominate at telomeres where the end-proximal Rap1 site is not too close from telomere double-stranded end. (**E**) At telomeres where the end-proximal Rap1 site is too close from the telomere double-stranded end to accommodate Ku when Rap1 is present, Rap1-Ku mutual exclusion predominates (same colour code as in panel D).

As shown in **Fig. 4B**, the frequency of Rap1 sites varied with the distance from the fusion points, displaying a notable peak at 2-4 bp and a distinct decrease at 8-25 bp. A similar pattern was observed in telomere fusions sequenced directly from genomic DNA, without the PCR amplification step (**Fig. S21C-D**). Rap1 depletion, which increases the frequency of telomere fusions ^33^ (**Fig. S21B**), led to a more uniform distribution of the Rap1 sites (**Fig. 4B**), indicating that Rap1 was responsible for the distinctive pattern observed in fusions from Rap1-proficient cells. To confirm the origin of these Rap1-dependent biases, we sequenced unfused telomeres from the same cells and found that the distribution of Rap1 sites remained relatively uniform up to the telomere ends (**Fig. 4C**). This shows that the large variations in Rap1 site frequency seen in fusions from Rap1-proficient cells were not present among the bulk of telomeres. Therefore, these variations must have resulted from a selection process favouring the fusion of telomeres whose end-proximal Rap1 sites were either very close to the end or more than ∼25-30 bp from the end.

As previously reported ^48^, NHEJ-dependent telomere fusion largely relies on Pol4, an NHEJ-specific DNA polymerase and the yeast orthologue of mammalian Polµ and Polλ (**Fig. S22A**). Interestingly, in the absence of Pol4, we observed that the pattern of Rap1 site frequency shifted by 1 bp closer to the fusion point (**Fig. S22B-C**). This finding is consistent with Pol4 filling-in the telomeric 3’ single-strand overhang by an average of 1 bp before end ligation. Consequently, fusions with a Rap1 site located 2 to 4 bp from the fusion point likely originated from telomeres where the end-proximal Rap1 site was initially only 1 to 3 bp away from the double-stranded telomeric end. Given that these telomeres are more prone to fusion, we conclude that the optimal positioning of Rap1 for end protection is crucial, as it aligns with its ability to accommodate Ku binding and effectively prevent the Ku inward translocation required for NHEJ (∼25 bp ^15^). This supports the idea that Rap1’s regulatory impact on Ku plays a fundamental role in shielding telomeres from fusion and underscores the significance of this mechanism for maintaining genome integrity *in vivo*.

## DISCUSSION

Preventing chromosome end fusion in cells where broken ends are rapidly sealed is a critical function of telomeres. Its efficiency depends on the synergistic action of multiple pathways that together reduce the likelihood of fusion to a very low level over the cell’s lifetime ^4 30 33 41 42^. We reveal that one of these pathways operates by directly modulating Ku’s binding mode at telomeres. This mechanism enables Ku to protect telomere ends from resection without engaging in NHEJ, a functional switch explaining Ku’s paradoxical presence at telomeres. By relying on a single, end-proximal telomere protein bound to DNA, the efficiency of this capping mechanism is decoupled from telomere length and the number of telomere-bound proteins, with just one being sufficient (**Fig. 4D**).

At double-strand breaks, Ku high-affinity for DNA ends can help it to overcome DNA-bound proteins that might otherwise impede NHEJ repair. However, this strong binding could pose a problem at telomeres. There, a control of Ku requires proteins with similar affinities for DNA. Remarkably, *S. cerevisiae* Rap1 meets this requirement ^12 34^ (**Fig. S3**). Our findings show that Rap1 can inhibit Ku NHEJ function at telomeres through distinct mechanisms, with their relative importance determined by Rap1 proximity to the telomere end (**Fig. 4D-E**). For a majority of telomeres, this distance is sufficient for Ku to bind without necessarily competing with Rap1 (**Fig. 4D**). In such cases, Rap1 acts as a “doorstop” restricting Ku’s inward translocation. This can leave it in an intermediate loading state where the DNA is not fully engaged in Ku’s binding site and the DNA end partially buried within the complex. This configuration prevents access of the DNA end to the catalytic domain of Lig4 ^15^ and to other enzymes required for NHEJ (e.g., Pol4), thereby inhibiting the subsequent ligation with another end. By limiting Ku inward translocation, Rap1 may also hinder the synapsis of ends ^15^, further contributing to NHEJ inhibition.

For a subset of telomeres, however, Rap1 is located too close to the end for functional cooperation with Ku, as the limited DNA accessible for Ku binding restricts its association with Rap1-bound telomere ends. As a result, Rap1 and Ku binding tend to become mutually exclusive (**Fig. 4E**). While this exclusion of Ku by Rap1 still prevents NHEJ, our sequencing of telomere fusions (**Fig. 4B**) indicates that this competitive mechanism is less effective than the one which targets Ku’s inward translocation, all else being equal. The vulnerability of this competition between Rap1 and Ku lies in the fact that when Rap1 temporarily dissociates, Ku can bind to the exposed DNA, translocate inward unimpeded, and initiate NHEJ. Although stronger Rap1 binding affinity might mitigate this limitation, it could come at the expense of making replication through telomeres more difficult ^49^, reflecting an evolutionary trade-off between efficient telomere protection and replication. Additionally, the most end-proximal Rap1 consensus site is sometimes positioned too far from the end to meaningfully impede Ku’s inward translocation and subsequent NHEJ. At these telomeres, the space between the last consensus Rap1 site and the telomere end is often occupied by alternative non-consensus Rap1 site (**Fig. S5**). Since these alternative sites are less efficient at inhibiting NHEJ (**Fig. S4**), telomeres with such configurations are less well protected.

These inherent limitations are offset by the presence of additional mechanisms that block NHEJ at telomeres, each with its own potential weaknesses. For instance, Rif2-dependent protection relies on specific protein interactions at telomere tips ^37 38^, which may not occur uniformly across all telomeres at all times. Crucially, telomere stability is maintained as long as at least one mechanism successfully prevent NHEJ at any given telomere at any given time. In wild-type yeast cells, telomere fusions are exceedingly rare (∼ 10^-6^) ^4^, underscoring the extent to which evolution has finely tuned telomere protection to ensure chromosome stability through the coordinated action of multiple overlapping pathways.

Our structure of the Rap1-DNA-Ku complex reveals that the co-binding of Rap1 and Ku on the same DNA molecule primarily depends on their respective protein-DNA interactions, with DNA acting as a hub for the formation of the ternary complex. Several factors may partially compensate for the lack of a significant specific interaction between Ku and Rap1. (1) Both Rap1 and Ku restrict transient strand separation at the DNA end, which may enhance their respective binding on DNA and contribute to the stability of the complex. (2) The configuration of Rap1 and Ku on DNA may reduce access to counter-ions, further stabilising their individual protein-DNA interactions. (3) When Ku binds to the Rap1-DNA complex, it contacts the C-terminal wrapping loop of Rap1, effectively locking this loop onto the DNA (patch 2, **Fig. S19**), thus reinforcing the stability of the Rap1-DNA association rather than antagonising it. This also suggests that the dynamics of the ternary complex relies on Ku’s ability to freely bind to and dissociate from Rap1-bound DNA ends.

These features can collectively compensate for the reduced Ku-DNA interface resulting from Rap1’s proximity to the DNA end, as well as the lack or weakness of a direct association between Ku and Rap1. However, this lack of a strong interaction may also reflect a critical aspect of this capping mechanism. A simulation based the law of mass action shows that even a 100-fold lower affinity of Ku for Rap1-bound DNA ends compared to free DNA ends can be sufficient to preserve the ternary complex, thereby protecting DNA ends from NHEJ (**Fig. S23**). A limit on the affinity might be evolutionarily advantageous, as a significant interaction between Ku and Rap1 could excessively seal the telomere ends, potentially hindering their replication. Overall, our findings highlight the robustness of a mechanism grounded in a fundamental property of these two proteins: their ability to bind tightly and independently their respective DNA substrate. This minimalistic design underscores the balance between DNA end protection and the flexibility required for telomere maintenance.

The remarkable fit between Rap1 and Ku on DNA implies that the shape compatibility of these two proteins might be under positive selection, optimising the minimal DNA length required downstream of Rap1 for Ku binding. This adaptation could enhance their co-binding at telomeres and thus strengthen the inhibition of NHEJ. Previous works hypothesised that a control of Ku’s inward translocation could take place at human and plant telomeres ^23 29^, suggesting a broader conservation of this mechanism. Additionally, the binding of telomeric DNA by tandems of Myb-like domains, which engage the major groove of DNA, is a conserved feature of telomere-associated proteins in many eukaryotes. This conservation may facilitate co-binding with Ku, as observed in yeast. The mechanism we propose could operate alongside other pathways inhibiting Ku’s NHEJ activity via direct protein-protein interactions ^13 27 50 51^. Such a dual control system of Ku, with complementary pathways acting in synergy, would be expected to ensure long term telomere stability.

## MATERIALS AND METHODS

See supplementary information.

## ACKNOWLEDGMENTS

The authors would like to thank Francisca Lottersberger, Daniela Rhodes, Sébastien Britton, Zhou Xu, Karine Dubrana, Pablo Radicella, and Thomas Guérin for fruitful discussions and comments; Claude Gazin and Jean-François Deleuze for support; Gérard Pehau-Arnaudet of Institut Pasteur (Paris) for his help to determine the conditions of cryofixation and for sample screening (detector financed by the Equipex CACSICE of the UBI platform); the European Synchrotron Radiation Facility for provision of beam time on CM01 and E. Kandiah for assistance; and the Molecular Biophysics facility (PFBMI) for access to mass photometry. Calculations were achieved on the HPC GPU resources made available by GENCI at TGCC@CEA (Joliot-Curie/Irene; allocation A0120313408) and at IDRIS@CNRS (Jean Zay; allocation AD010313694). Research in the laboratory of S.Marc. was supported by grants from *Fondation pour la Recherche Médicale* (EQU202203014702), *Agence Nationale de la Recherche* (ANR-15CE12-0007, ANR-22-CE12-0037), *Fondation ARC pour la Recherche sur le Cancer*, *Ligue contre le Cancer*, CEA Radiation biology program, GGP CEA EDF program and *Inserm*. JBC, PFV and PC thank support from *Agence Nationale de la Recherche* (ANR-22-CE12-0037, ANR 23-CE11-0033), FRISBI ANR-10-INBS-0005, CEA Radiation biology program and GGP CEA EDF program. ELC, GM and SB thank support from *Agence Nationale de la Recherche* (ANR-22-CE12-0037), *Fondation ARC pour la Recherche sur le Cancer* and *ERM-Université Paris-Saclay*. This work is dedicated to Armando Felsani, a brilliant scientist and great mentor who passed away on 21 April 2022.

## DATA AND MATERIALS AVAILABILITY

The atomic coordinates of the ternary complex Rap1-DNA-Ku and the binary complex DNA-Ku have been deposited in the Worldwide Protein Data Bank (wwPDB) with accession codes 8S8P and 8S82, respectively. The corresponding maps have been deposited in the Electron Microscopy Data Bank (EMDB) with accession codes EMD-19811 and EMD-19790, respectively.

## REFERENCES

1. de Lange, T. Shelterin-Mediated Telomere Protection. Annu. Rev. Genet. 52, 223–247 (2018).

2. Ruis, P. & Boulton, S. J. The end protection problem—an unexpected twist in the tail. Genes Dev. 35, 1–21 (2021).

3. Chan, S. W.-L. & Blackburn, E. H. Telomerase and ATM/Tel1p Protect Telomeres from Nonhomologous End Joining. Molecular Cell 11, 1379–1387 (2003).

4. Pobiega, S., Alibert, O. & Marcand, S. A new assay capturing chromosome fusions shows a protection trade-off at telomeres and NHEJ vulnerability to low-density ionizing radiation. Nucleic Acids Research 49, 6817–6831 (2021).

5. Stuart, A. & De Lange, T. Replicative senescence is ATM driven, reversible, and accelerated by hyperactivation of ATM at normoxia. Preprint at 10.1101/2024.06.24.600514 (2024).

6. Marcand, S., Gilson, E. & Shore, D. A Protein-Counting Mechanism for Telomere Length Regulation in Yeast. Science 275, 986–990 (1997).

7. Teixeira, M. T., Arneric, M., Sperisen, P. & Lingner, J. Telomere Length Homeostasis Is Achieved via a Switch between Telomerase-Extendible and -Nonextendible States. Cell 117, 323–335 (2004).

8. Krauskopf, A. & Blackburn, E. H. Control of telomere growth by interactions of RAP1 with the most distal telomeric repeats. Nature 383, 354–357 (1996).

9. Tesmer, V. M., Brenner, K. A. & Nandakumar, J. Human POT1 protects the telomeric ds-ss DNA junction by capping the 5′ end of the chromosome. Science 381, 771–778 (2023).

10. Zahid, S. et al. The Multifaceted Roles of Ku70/80. IJMS 22, 4134 (2021).

11. Walker, J. R., Corpina, R. A. & Goldberg, J. Structure of the Ku heterodimer bound to DNA and its implications for double-strand break repair. Nature 412, 607–614 (2001).

12. Pfingsten, J. S. et al. Mutually Exclusive Binding of Telomerase RNA and DNA by Ku Alters Telomerase Recruitment Model. Cell 148, 922–932 (2012).

13. Chen, H. et al. Structural Insights into Yeast Telomerase Recruitment to Telomeres. Cell 172, 331–343.e13 (2018).

14. Nemoz, C. et al. XLF and APLF bind Ku80 at two remote sites to ensure DNA repair by non-homologous end joining. Nat Struct Mol Biol 25, 971–980 (2018).

15. Chen, S. et al. Structural basis of long-range to short-range synaptic transition in NHEJ. Nature 593, 294–298 (2021).

16. Palmbos, P. L., Daley, J. M. & Wilson, T. E. Mutations of the Yku80 C Terminus and Xrs2 FHA Domain Specifically Block Yeast Nonhomologous End Joining. Mol Cell Biol 25, 10782–10790 (2005).

17. Graham, T. G. W., Walter, J. C. & Loparo, J. J. Two-Stage Synapsis of DNA Ends during Non-homologous End Joining. Molecular Cell 61, 850–858 (2016).

18. Chaplin, A. K. et al. Cryo-EM of NHEJ supercomplexes provides insights into DNA repair. Molecular Cell 81, 3400–3409.e3 (2021).

19. Gravel, S., Larrivée, M., Labrecque, P. & Wellinger, R. J. Yeast Ku as a Regulator of Chromosomal DNA End Structure. Science 280, 741–744 (1998).

20. Baumann, P. & Cech, T. R. Protection of Telomeres by the Ku Protein in Fission Yeast. MBoC 11, 3265–3275 (2000).

21. d’Adda Di Fagagna, F., et al. Effects of DNA nonhomologous end-joining factors on telomere length and chromosomal stability in mammalian cells. Current Biology 11, 1192–1196 (2001).

22. Wang, Y., Ghosh, G. & Hendrickson, E. A. Ku86 represses lethal telomere deletion events in human somatic cells. Proc. Natl. Acad. Sci. U.S.A. 106, 12430–12435 (2009).

23. Bombarde, O. et al. TRF2/RAP1 and DNA–PK mediate a double protection against joining at telomeric ends. EMBO J 29, 1573–1584 (2010).

24. Vodenicharov, M. D., Laterreur, N. & Wellinger, R. J. Telomere capping in non-dividing yeast cells requires Yku and Rap1. EMBO J 29, 3007–3019 (2010).

25. Lopez, C. R. et al. Ku Must Load Directly onto the Chromosome End in Order to Mediate Its Telomeric Functions. PLoS Genet 7, e1002233 (2011).

26. Sfeir, A. & De Lange, T. Removal of Shelterin Reveals the Telomere End-Protection Problem. Science 336, 593–597 (2012).

27. Ribes-Zamora, A., Indiviglio, S. M., Mihalek, I., Williams, C. L. & Bertuch, A. A. TRF2 Interaction with Ku Heterotetramerization Interface Gives Insight into c-NHEJ Prevention at Human Telomeres. Cell Reports 5, 194–206 (2013).

28. Mateos-Gomez, P. A. et al. Mammalian polymerase θ promotes alternative NHEJ and suppresses recombination. Nature 518, 254–257 (2015).

29. Valuchova, S. et al. Protection of Arabidopsis Blunt-Ended Telomeres Is Mediated by a Physical Association with the Ku Heterodimer. Plant Cell 29, 1533–1545 (2017).

30. Myler, L. R. et al. DNA-PK and the TRF2 iDDR inhibit MRN-initiated resection at leading-end telomeres. Nat Struct Mol Biol 30, 1346–1356 (2023).

31. Wellinger, R. J. & Zakian, V. A. Everything You Ever Wanted to Know About *Saccharomyces cerevisiae* Telomeres: Beginning to End. Genetics 191, 1073–1105 (2012).

32. König, P., Giraldo, R., Chapman, L. & Rhodes, D. The Crystal Structure of the DNA-Binding Domain of Yeast RAP1 in Complex with Telomeric DNA. Cell 85, 125–136 (1996).

33. Marcand, S., Pardo, B., Gratias, A., Cahun, S. & Callebaut, I. Multiple pathways inhibit NHEJ at telomeres. Genes Dev. 22, 1153–1158 (2008).

34. Williams, T. L., Levy, D. L., Maki-Yonekura, S., Yonekura, K. & Blackburn, E. H. Characterization of the Yeast Telomere Nucleoprotein Core. Journal of Biological Chemistry 285, 35814–35824 (2010).

35. Pobiega, S. & Marcand, S. Dicentric breakage at telomere fusions. Genes Dev. 24, 720–733 (2010).

36. Lescasse, R., Pobiega, S., Callebaut, I. & Marcand, S. End-joining inhibition at telomeres requires the translocase and polySUMO-dependent ubiquitin ligase Uls1. EMBO J 32, 805–815 (2013).

37. Roisné-Hamelin, F. et al. Mechanism of MRX inhibition by Rif2 at telomeres. Nat Commun 12, 2763 (2021).

38. Khayat, F. et al. Inhibition of MRN activity by a telomere protein motif. Nat Commun 12, 3856 (2021).

39. Bae, N. S. & Baumann, P. A RAP1/TRF2 Complex Inhibits Nonhomologous End-Joining at Human Telomeric DNA Ends. Molecular Cell 26, 323–334 (2007).

40. Sarthy, J., Bae, N. S., Scrafford, J. & Baumann, P. Human RAP1 inhibits non-homologous end joining at telomeres. EMBO J 28, 3390–3399 (2009).

41. Okamoto, K. et al. A two-step mechanism for TRF2-mediated chromosome-end protection. Nature 494, 502–505 (2013).

42. Benarroch-Popivker, D. et al. TRF2-Mediated Control of Telomere DNA Topology as a Mechanism for Chromosome-End Protection. Molecular Cell 61, 274–286 (2016).

43. Khayat, F., Alshmery, M., Pal, M., Oliver, A. W. & Bianchi, A. Binding of the TRF2 iDDR motif to RAD50 highlights a convergent evolutionary strategy to inactivate MRN at telomeres. Nucleic Acids Research gkae509 (2024) doi:10.1093/nar/gkae509.

44. Buchman, A. R., Kimmerly, W. J., Rine, J. & Kornberg, R. D. Two DNA-Binding Factors Recognize Specific Sequences at Silencers, Upstream Activating Sequences, Autonomously Replicating Sequences, and Telomeres in *Saccharomyces cerevisiae*. Molecular and Cellular Biology 8, 210–225 (1988).

45. Analikwu, B. T., et al. Telomere Protein Arrays Stall DNA Loop Extrusion by Condensin. http://biorxiv.org/lookup/doi/10.1101/2023.10.29.564563 (2023) doi:10.1101/2023.10.29.564563.

46. Bonetti, D. et al. DNA binding modes influence Rap1 activity in the regulation of telomere length and MRX functions at DNA ends. Nucleic Acids Research 48, 2424–2441 (2020).

47. Matot, B. et al. The orientation of the C-terminal domain of the Saccharomyces cerevisiae Rap1 protein is determined by its binding to DNA. Nucleic Acids Research 40, 3197–3207 (2012).

48. Pardo, B., Ma, E. & Marcand, S. Mismatch Tolerance by DNA Polymerase Pol4 in the Course of Nonhomologous End Joining in *Saccharomyces cerevisiae*. Genetics 172, 2689–2694 (2006).

49. Douglas, M. E. & Diffley, J. F. X. Budding yeast Rap1, but not telomeric DNA, is inhibitory for multiple stages of DNA replication in vitro. Nucleic Acids Research 49, 5671–5683 (2021).

50. Taddei, A., Hediger, F., Neumann, F. R., Bauer, C. & Gasser, S. M. Separation of silencing from perinuclear anchoring functions in yeast Ku80, Sir4 and Esc1 proteins. EMBO J 23, 1301–1312 (2004).

51. Rai, R. et al. Homology directed telomere clustering, ultrabright telomere formation and nuclear envelope rupture in cells lacking TRF2B and RAP1. Nat Commun 14, 2144 (2023).

52. Forstemann, K., Höss, M. & Lingner, J. Telomerase-dependent repeat divergence at the 3’ ends of yeast telomeres. Nucleic Acids Research 28, 2690–2694 (2000).

