## Supplementary material for "Restriction on Ku’s Inward Translocation Caps Telomere Ends": SI

### MATERIALS AND METHODS

#### Yeast strains and molecular genetics

Strains used in this study are listed in **Table S3**. Survival to I-SceI cleavage was determined as previously described (Roisné-Hamelin et al. 2021). Briefly, cells were grown to saturation for 16 hrs in synthetic medium lacking uracil supplemented with glucose (2%). The cells were then diluted in water and spread onto synthetic medium plates containing galactose (2%). Colonies were counted after 4 days at 30°C.

#### Chromosome Fusion Capture assay (CFC)

To determine survival to *CEN6* loss, cells grown on rich glucose medium (YPD) plates are picked and plated on glucose medium plates lacking uracil to counter-select rare cells that had already lost *CEN6* due to Cre background expression in the absence of galactose. After 24 hrs of growth at 30°C, distinct re-streaks were used to inoculate separate liquid cultures in YPD. Each culture was then grown to saturation for 3 days at 30°C before plating on a galactose medium plate lacking leucine ($5 \times 10^7$ cells per plate or fewer if the frequency of survival to *CEN6* loss is above $4 \times 10^{-6}$). Colonies were counted after 5 to 7 days at 30°C. Each data point is derived from an independent re-streak and an independent cell culture spread on a single plate.

#### ChIP assay

The Chip assay was performed as previously described (Hafner et al. 2018). 40 mL of G1-arrested cells (treatment with alpha-factor $10^{-7}$ M, approximately $10^7$ cells/mL) were fixed in 1% formaldehyde for 10 min while rotating and then quenched with 2 mL 2.5 M glycine. Next, the cells were washed with ice cold HBS (50 mM HEPES pH 7.8, 140 mM NaCl), and frozen in 600 µL of ChIP lysis buffer (50 mM HEPES pH 7.8, 140 mM NaCl, 1 mM EDTA, 1% NP40, 1% sodium deoxycholate) complemented with protease inhibitors (Roche 04693132001). Cells were lysed using a FastPrep (MP Biomedicals), 3x30 seconds at 6 M/s and zirconium/silica beads (BioSpec Products 11079105z). The beads were then removed and cells were sonicated at a high setting, with 30 sec on 30 sec off for 3 min using a Diagenode sonicator at 4°C. Sonicates were clarified by spinning at top speed for 25 min at 4°C. 10 µl of solution were incubated at 65°C for 16 h (input samples, to normalize the qPCR step). Immunoprecipitation was performed for 1 hr at 4°C with a primary antibody (anti-Myc 9E10 from ThermoFisher Scientific or anti-Rap1, a gift from David Shore laboratory) followed by 2 hrs at 4°C with beads coupled to a secondary antibody (sheep anti-mouse or sheep anti-rabbit from Invitrogen, references 11201D and 11203D respectively). For washes, samples were placed on a magnet to collect beads. Beads were first rinsed in 1 mL ChIP lysis buffer and then washed with AT1 (50 mM HEPES pH 7.8, 140 mM NaCl, 1mM EDTA, 0.03% SDS), AT2 (50 mM HEPES pH 7.8, 1M NaCl, 1 mM EDTA), AT3 (20 mM Tris-HCl pH 7.5, 250 mM LiCl, 1 mM EDTA, 0.5% NP40, 0.5% sodium deoxycholate) and AT4 (20 mM Tris-HCl pH 7.5, 0.1 mM EDTA) wash buffers. Samples were then eluted in 140 µL of elution buffer (AT4 + 1% SDS) and the crosslink reverted in 140 µL of elution buffer at 65°C for 16 hrs. Subsequently, samples were digested with proteinase K in TE buffer for 1 hour at 37°C and then cleaned-up using a QIAquick PCR Purification Kit (Qiagen 28104). qPCRs were performed in a 20 µL volume with PowerSYBR green PCR master mix (Applied Biosystems 4367659) and an Applied Biosystems HT7900 machine.

**Telomere and telomere fusions sequencing**

Fused and unfused telomeres were amplified by PCR as described previously (Lescasse et al. 2013). PCR products obtained from three independent cell cultures were pooled and sequenced by Nanopore sequencing. Base calling was achieved with the ONT Guppy software in super accurate mode (configured with the dna_r9.4.1_450bps_sup.cfg file). We used the cpu version 6.0.1 of Guppy on a computing cluster. Using Seqkit v2.1.0 (Shen et al. 2016), we mapped telomere sequences matching the motif 5'$G_{0-3}(TG_{1-3})_{\geq 3}[N_{0-2}(TG_{1-3})_{\geq 3}]_{\geq 1}T_{0-1}$3' (*i.e.* up to 2 divergent bases every 3 $TG_{1-3}$ repeats). The fusion points were determined where a reverse C-rich telomere sequence directly follows a forward G-rich telomere sequence. The 3' ends of unfused telomeres were positioned where a polyC stretch directly follows a forward G-rich telomere sequence. We used Seqkit to search for the Rap1 sites. The distance to the fusion point refers to the distance of the fusion-proximal base of the Rap1 site, while the distance to the telomere 3' end refers to the distance of the last base before the polyC stretch. We used dedicated *Perl* scripts to generate the Rap1 site frequency maps. Due to sequencing errors, some sites present in the original sequences were missed. With a lower probability, some were created. The sequencing of the telomere G-rich strand proved to be more accurate than sequencing of the telomere C-rich strand, *i.e.* the telomere G-rich strands display fewer divergent bases in the $TG_{1-3}$ repeats, resulting in higher frequencies of Rap1 site.

**Protein purification**

The 6His-$Rap1_{[1-827]}$ construct was cloned into the pETM-13 vector and protein expression was induced with 0.5 mM isopropyl-ß-D- thiogalactoside (IPTG) at 20 °C overnight into *E. coli* strain BL21 (DE3) STAR. All of the subsequent protein purification steps were carried out at 4 °C. Cells were harvested, suspended in lysis buffer (20 mM Tris-HCl pH 8.0, 200 mM NaCl, 1 mM DTT, 1 mg/mL lysozyme, 1 mM 4-(2-aminoethyl) benzenesulphonyl fluoride (AEBSF), 10 mM benzaminide, 2 µM pepstatin) and disrupted by sonication. The extract was cleared by centrifugation at 186,000 g for 1 hr and then incubated with NiNTA resin (Qiagen) for 3 hrs. The mixture was poured into an Econo-Column® Chromatography column (Biorad). After extensive washing of the resin with buffer A (20 mM Tris-HCl pH 8.0, 1 M NaCl, 1 mM DTT, 20 mM imidazole) and then with buffer B (20 mM Tris-HCl pH 8.0, 50 mM NaCl, 1 mM DTT) complemented with 20 mM imidazole, the protein was eluted with buffer B complemented with 400 mM imidazole. Fractions containing purified 6His-$ScRap1_{[1-827]}$ were pooled and applied to a 1-ml Resource Q column (Cytiva) equilibrated with buffer B. After, a linear gradient of NaCl from 50 to 600 mM was applied. The fractions having the 6His-$Rap1_{[1-827]}$ were then directly loaded onto a 1ml HiTrap Heparin column (Cytiva) equilibrated with buffer B and a linear gradient of NaCl from 100 to 1000 mM was applied. Purified 6His-Rap1$_{[1-827]}$ was stored at −80 °C.

*S. cerevisiae* Ku was expressed in insect cells (sf21) with a 10His tag on the Yku80 N-terminus. The pellets were suspended in lysis buffer (20 mM MES pH 6.0, 850 mM NaCl, 150 mM KCl, 1 mM DTT, 40 mM imidazole, 10% glycerol, ½ tablet of protease inhibitor cocktail (for 250 mL) (cOmplete EDTA free, Roche diagnostics) and sonicated in 3 cycles of 1 min at 60% amplitude (Vibra cell VC750; Sonics & Materials inc.). Then, 2 µL of benzonase (0.01 U/µl) and 10 mM MgCl were added and the lysate was incubated for 20 min followed by a centrifugation step (20,000 rpm for 20 min). All the previous steps were performed at temperature lower than 15 °C. The obtained supernatant was loaded on a NiNTA column (20 mL, pre-equilibrated with the lysis buffer) and incubated for 1h at 4 °C under agitation. The column was washed 3 times with 50 mL of lysis buffer and the protein was eluted using an elution buffer (20 mM MES pH 6.0, 300 mM NaCl, 300 mM imidazole, 1 mM DTT). The elution fractions were dialyzed against QA buffer (20 mM MES pH 6.0, 50 mM KCl, 50 mM NaCl, 1 mM DTT, 2 mM EDTA) at 4 °C using 6-8 kDa membranes (SpectraPor). The dialyzed sample was then injected onto a ResQ column (6 mL from Cytivia). The protein of interest was eluted with a salt gradient. The fractions of the ResQ were dialyzed against 1 L of storage buffer (20 mM MES pH 6.0, 500 mM NaCl, 5 % glycerol, 1 mM DTT) for 16 hrs at 4 °C and frozen in liquid nitrogen.

**Electrophoresis mobility shift assay (EMSA)**

The protein-DNA binding reactions were conducted by incubating the purified proteins with DNA duplexes labelled with Cy5 at the 5' end. The incubation was performed in hybridization buffer consisting of 20 mM Tris-HCl (pH 8.0), 150 mM NaCl, 1 mM DTT, 10% glycerol, 5 mM MgCl2 and 0.3 mg/ml BSA. The total reaction volume was 10 µl, with a final duplex DNA concentration of 1 nM. Incubation was carried out at 25°C from 2 to 16 hrs. The extended incubation time ensured that the binding equilibrium between the proteins and DNA was reached (Pfingsten et al. 2012). The protein-DNA complexes were resolved by electrophoresis using a vertical 6.7% acrylamide gel prepared with a ratio of acrylamide to bis-acrylamide of 19:1 (Biorad #1610144) in glycerol-based TBE buffer. Electrophoresis was conducted at 220 V for 3.5 hrs to separate the protein-DNA complexes based on their mobility shifts. Gel images were acquired by scanning the gels using a Typhoon imager.

**Mass photometry**

Mass photometry is a label-free single molecule approach based on light scattering by single particles at a glass-water interface, allowing for an estimate of the sizes of molecular complexes (Young et al. 2018). Mass photometry data were acquired using a Refeyn TwoMP mass photometer located at the Institut Pasteur in Paris, France. Data acquisition was performed using the AcquireMP software. 18 µl of solution buffer (20 mM Tris-HCl, pH 7.5, 100 mM NaCl) was loaded onto the objective of the mass photometer, followed by the addition of 2 µl of the sample. The sample was allowed to incubate for 60 seconds on the objective. Bovine serum albumin (BSA) and urease were used as calibration standards. The DiscoverMP software was utilized to convert the measured values to molecular masses. Rap1 and Ku proteins were prepared at a final concentration of 50 nM. For experiments involving different DNA duplexes, Rap1 and Ku were mixed at a 1:1:1 ratio with the respective DNA duplexes and incubated together for 5 min prior to measurement. Duplicate measurements were performed for each experimental condition.

**SwitchSENSE**

All switchSENSE® experiments were performed on the heliX$^+$ biosensor using ADP-48-2-0 heliX Adapter chips (Dynamic Biosensors, Germany). The chip was regenerated with a high-pH regeneration solution (HK-REG-1, Dynamic Biosensors, Germany) and freshly functionalized with the target DNA-sequences (ella Biotech GmbH, Germany) hybridized to red adapters strands (AS-1-Ra, Dynamic Biosensors, Germany) before each measurement or concentration determination. The experiments were performed in TE140 buffer (10 mM Tris-HCl, 140 mM NaCl, 50 µM EDTA, 50 µM EGTA, 0.05% Tween20, pH 7.4) buffer. The kinetic interaction was tracked by fluorescence proximity sensing (FPS), where changes in the local environment of the reporter dye affect its fluorescence signal. A simultaneous measurement on a second detection spot without the DNA-target sequence was recorded and used as real-time control to monitor and subtract unspecific binding. In addition to this reference, a blank run (buffer only) was also recorded and subtracted to obtain the net interaction signal. The experiments were performed at a negative potential, a sampling rate of 1 Hz, a temperature of 25°C and the association and dissociation flowrates were 200 µL/min and 500 µL/min, respectively. For measurements with Ku protein, 10 µM of a single strand 20T oligo (IDT, Belgium) was added during association to suppress non-specific binding effects. Experiment design, workflow and data analysis were performed with the heliOS software (Dynamic Biosensors, Germany). All data was generated at least in triplicates and the results were averaged.

**Cryo-EM of Rap1-DNA-Ku complex**

Purified 6His-tagged Rap1$_{[1-827]}$ protein (6 µM) was incubated 10 min at room temperature with a 21 bp duplex DNA containing a Rap1 consensus site and a 5 bp downstream extension (3 µM). Subsequently, purified Ku protein (6 µM) was added to the solution, which was then further incubated for 10 min at room temperature in a buffer composed of 10 mM Tris-HCl (pH 8.0) and 50 mM NaCl. The freshly formed Rap1-DNA-Ku complex was subjected to purification by size exclusion chromatography using a Superose 6 Increase 3.2/300 column (Cytiva) and then concentrated using a 30 kDa molecular weight cut-off Amicon centrifugal filter unit.

Three microliters of the purified complex were applied onto glow-discharged holey carbon grids (Quantifoil Cu R1.2/1.3, 300 mesh). The grids were subsequently blotted for 3 sec with zero force and plunge-frozen in liquid ethane using a FEI Vitrobot Mark IV plunger (Thermo Fisher Scientific) set at 4°C and 95% humidity.

Cryo-electron microscopy data were acquired on a Titan Krios microscope equipped with a Gatan K3 camera at the European Synchrotron Radiation Facility in Grenoble (Kandiah et al. 2019). A total of 28,670 movies were recorded. All data collection parameters are provided in **Table S2**.

**Image processing and three-dimensional reconstruction**

The data were processed using cryoSPARC v4 software. After motion correction and CTF estimation, movies with a CTF fit resolution greater than 5.5 Å and ice thickness exceeding 1.2 Å were excluded, resulting in the selection of 24,181 micrographs. Subsequent blob picking identified a first set of 663,798 particles that were extracted with a box size of 512 pixels. Following 2D classification, 15 classes were chosen for template picking, resulting in the extraction of a second set of 13,650,337 particles. 1,544,386 particles were selected for 3D reconstruction after further 2D classification. Three-dimensional reconstructions of the ternary and binary complexes were obtained after *ab initio* reconstruction (3 classes) followed by heterogeneous refinement. Subsequent homogenous refinement followed by non-uniform refinement yielded to the final 3D maps with resolutions of 3.11 Å and 2.92 Å for the ternary and binary complexes, respectively. The overall image processing workflow is summarized schematically in **Sup. Fig. S12**.

**Molecular modelling**

For the two models, protein and nucleic chain preparation as well as initial model assembly were achieved with the *CHARMM v47b1* software (Brooks et al. 2009). All energy minimization and molecular dynamics calculations were performed in the *CHARMM36m* force field (Huang et al. 2017). The final structural model of the $Rap1_{DBD}$-DNA-Ku complex was obtained in two steps: first, we built a starting model that could fulfil the EMSA data obtained with different duplex lengths. This model was refined by molecular dynamics flexible fitting (MDFF) and finally optimized by real space refinement with *PHENIX* (v.1.21-rc1-49.58). To construct the initial molecular model of the $Rap1_{DBD}$-DNA-Ku complex we used the X-ray structure of the *S. cerevisiae* $Rap1_{DBD}$ domain in complex with a DNA duplex (PDB:3UKG) (Matot et al. 2012). We modified the DNA structure in 3UKG to fit the exact sequence of the 21-mer used in the present study (the 5'-GTGGTGTGTGGGTGTGTGTGT-3' G-rich strand and the complementary 5'-ACACACACACCCACACACCAC-3' C-rich strand) using the *swapna* command of *ChimeraX 1.6.1* (Goddard et al. 2018, Pettersen et al. 2021). This DNA sequence corresponds to the minimal length duplex determined *in vitro* for simultaneous Ku/Rap1 binding. In the 3UKG crystal structure, some loops of the $Rap1_{DBD}$, which are not directly involved in interactions with the DNA duplex, are missing. These loops were rebuilt using the loop modelling protocol of *Modeller 10.4* (Šali and Blundell 1993, Eswar 2003). We also selected the *S. cerevisiae* Yku70/Yku80 complex in interaction with a telomerase *TCL1* RNA fragment (PDB:5Y58 (Chen et al. 2018)). We used the *TLC1* RNA as a template to set the position of the 21-mer-DNA-$Rap1_{DBD}$ complex with respect to Ku. Step-by-step translation-rotation, nucleotide by nucleotide, was used to preserve the interactions between the 21-mer DNA phosphate backbone and Ku at each step. These motions were continued until

$Rap1_{DBD}$ reached the closest possible position to Ku without any significant steric clash between the two partners. The resulting model was inspected graphically and some residual minor clashes were manually corrected. We then saved the cartesian coordinates of Yku70, Yku80, the 21-mer DNA duplex and $Rap1_{DBD}$ to form the initial complex. This initial model was fitted in the cryo-EM map using the *Volume* Tool of *ChimeraX*. The initial DNA-Ku model was obtained by fitting Ku and the DNA duplex separately in the cryo-EM map corresponding to the binary complex.

Next, the refinement of the ternary $Rap1_{DBD}$-DNA-Ku complex and the binary DNA-Ku complex under the restraint of the cryo-EM map was achieved using molecular dynamics flexible fitting (MDFF), in explicit water using the *NAMD* software (Phillips et al. 2005) followed by a final step of real-space refinement using the RealSpaceRefine protocol of the Phenix Software (Liebschner et al. 2019). The MDFF protocol in the presence of explicit water (Trabuco et al. 2008) was used because this approach led to significantly better results than the *in vacuo* one, particularly regarding the quality of protein geometry (data not shown). For each complex, the model was immersed in a water box with size chosen such that there is at least a 12 Å water layer between the solute and the edge of the water box in the x, y and z directions using the *solvate* plugin of the *VMD* software (Humphrey et al. 1996). The system was then neutralized, and the NaCl concentration set to 150 mM using the *autoionize* plugin of *VMD*. The main steps of the MDFF protocol are described below. The original cryo-EM map in *mrc* format was trimmed around the atoms of the initial model using the *Voledit* program of the *Situs* suite (Wriggers & Chacón. 2001) ensuring that the resulting map was entirely included in the water box. Then, the trimmed map was converted to *dx* format and transformed into a grid potential readable by *NAMD* using the *mdff* utility of *VMD* (Trabuco et al. 2008). A set of restraints were defined to preserve a correct geometry of the proteins and 21-mer DNA as the grid potential could exert significant forces that may alter the correct geometry of the simulated partners, including the correct chirality of some atoms and the configuration of the peptide planes. This was achieved using the *ssrestraints* plugin of VMD. The NAMD protocol started with 20000 steps energy minimization during which the system was restrained by the grid derived from the cryo-EM map with a scaling factor of 0.3 along with the geometrical restraints described above. This energy minimization was followed by 10 ns *MD* simulation (scaling factor 0.3). During this simulation, the map restraint energy was monitored. At 4 ns, this energy reached a stable minimum, indicating that the length of the simulation was sufficient to refine the structure with this protocol. Finally, 30,000 steps of energy minimization were calculated with the geometrical restraints and the grid potential with a scale factor of 10 on the latter term.

At last, the model resulting from the MDFF protocol was submitted to a refinement step using the *RealSpaceRefine* protocol of *Phenix* (v.1.21-rc1-49.58) (Afonine et al. 2018). This protocol includes systematic search for bad rotamers with respect to backbone dependent conformation rotamer database and poor agreement with the cryo-EM map. Secondary structure restraints defined for protein structure (alpha-helices and beta-strand structures), Ramachandran potential and nucleic acid secondary structure restraints (inter-strand base paring and intra-strand base stacking) were also used during energy minimization and molecular dynamics to maintain a correct geometry of the model during refinement. It employs 5 cycles of energy minimization and simulated annealing (from 5000 K to 300 K) with the above restraints to improve the fit of the cryo-EM map while producing a higher-quality structure than the MDFF protocol alone. Local resolution maps were calculated using the two half electron density maps obtained for the $Rap1_{DBD}$-DNA-Ku ternary complex and the DNA-Ku binary complex using the *LocalResolutionMap* protocol of *Phenix* (v.1.21-rc1-49.58). All MD trajectories were calculated with the *NAMD* 2.14 CUDA version on the HPC GPU resources made available by GENCI at TGCC@CEA (Joliot-Curie/Irene; allocation A0120313408) and at IDRIS@CNRS (Jean Zay; allocation AD010313694).

**Equilibrium Binding Simulations:**

The equilibrium binding simulations were achieved with the program DynaFit4 (Kuzmic 1996, Kuzmic 2006). This software, originally developed for statistical analysis of experimental data in biochemistry and to simulate enzyme kinetics and inhibition, is based on the law of mass action. It allows for the representation of the equilibria using a straightforward syntax. The number of equilibria is, at least in principle, unlimited, as are the numbers of reactants and products. Each equilibrium can be represented by a dissociation (or association) constant, as well as by kinetics constant. The rigorous solution of such large systems is facilitated by the generalized solver implemented in DynaFit4, which efficiently solves the differential equations beside the simple syntax describing the different equilibria. DynaFit4 does not make any assumption regarding the concentration ratio. Users can set values for every initial concentration and $k_{on}/k_{off}$ for each equilibrium matching the experimental values to fit a model to a set of experimental data or allow the solver to optimize certain values for data fitting. It is also possible to simulate data for evaluating the impact of any parameter (initial concentrations, $k_{on}/k_{off}$, $K_d$) on the products. DynaFit4 can simulate progress curves (concentrations as function of time) as well as equilibrium data. In this work, progress curves were systematically calculated to evaluate the relative evolution of the concentrations of the reactants and products for various parameter sets, ensuring that the equilibrium values were reached. The progress curves were simulated from 0 to 3600 sec, which was sufficient for all concentration to reach its equilibrium value.

### SUPPLEMENTARY TABLES

**Table S1.** Statistical analysis of Fig. 1C, S4 and S20B
(*t* test unpaired two tails)

| **Figure 1C** | no insert *rif2Δ sir4Δ* | d11 *rif2Δ sir4Δ* |
|---|---|---|
| no insert WT | 0.51 NS | <0.0001 **** |
| d11 WT | <0.0001 **** | 0.64 NS |
| no insert *rif2Δ sir4Δ* | | <0.0001 **** |
| d41 *rif2Δ sir4Δ* | 0.0049 ** | <0.0001 **** |
| d31 *rif2Δ sir4Δ* | 0.16 NS | <0.0001 **** |
| d29 *rif2Δ sir4Δ* | 0.011 * | <0.0001 **** |
| d26 *rif2Δ sir4Δ* | <0.0001 **** | <0.0001 **** |
| d21 *rif2Δ sir4Δ* | <0.0001 **** | <0.0001 **** |
| d19 *rif2Δ sir4Δ* | <0.0001 **** | 0.0021 ** |
| d16 *rif2Δ sir4Δ* | <0.0001 **** | 0.85 NS |
| d13 *rif2Δ sir4Δ* | <0.0001 **** | 0.21 NS |
| d11 *rif2Δ sir4Δ* | <0.0001 **** | |
| mut1 | <0.0001 **** | <0.0001 **** |
| mut2 | 0.051 NS | <0.0001 **** |
| mut3 | 0.011 * | <0.0001 **** |
| random | 0.18 NS | <0.0001 **** |
| **Figure S4** | no insert | telomere site |
| no insert | | <0.0001 **** |
| telomere site | <0.0001 **** | |
| tel. alt. 1 | <0.0001 **** | <0.0001 **** |
| tel. alt. 2 | <0.0001 **** | <0.0001 **** |
| tel. alt. 3 | <0.0001 **** | <0.0001 **** |
| HMR-E | <0.0001 **** | <0.0001 **** |
| TEF2-II | <0.0001 **** | <0.0001 **** |
| RPS17 | <0.0001 **** | <0.0001 **** |
| RPS11B | 0.0099 ** | <0.0001 **** |
| RPL41-II | 0.10 NS | <0.0001 **** |
| **Figure S20B** | no insert *YKU80*$^{+}$ | Rap1 site *YKU80*$^{+}$ |
| no insert *YKU80*$^{+}$ | | <0.0001 **** |
| no insert *yku80-EE* | 0.018 * | <0.0001 **** |

| | | |
|---|---|---|
| no insert *yku80-AEE* | <0.0001 **** | <0.0001 **** |
| no insert *yku80-AAA* | 0.0010 *** | <0.0001 **** |
| no insert *yku80-EEEA* | <0.0001 **** | <0.0001 **** |
| Rap1 site *YKU80*+ | <0.0001 **** | |
| Rap1 site *yku80-EE* | <0.0001 **** | 0.23 NS |
| Rap1 site *yku80-AEE* | <0.0001 **** | <0.0001 **** |
| Rap1 site *yku80-AAA* | <0.0001 **** | <0.0001 **** |
| Rap1 site *yku80-EEEA* | <0.0001 **** | <0.0001 **** |

**Table S2. Cryo-EM data collection parameters and refinement statistics**

| | | |
|---|---|---|
| **Data collection and processing** | **$Rap1_{DBD}$-DNA-Ku and DNA-Ku** | |
| Microscope | Titan | |
| Detector | Gatan K3 | |
| Magnification | 130K | |
| Energy filter slit width (eV) | 20 eV | |
| Automation software | EPU | |
| Voltage (kV) | 300 | |
| Electron exposure (e–/Å$^2$) | 52 | |
| Defocus range (μm) | -0.75 to -2.5 in 0.25 μm step | |
| Pixel size (Å) | 0.657 | |
| Symmetry imposed | C1 | |
| Micrographs used | 24,181 | |
| Initial particle images (no.) | 1,544,386 | |
| | **$Rap1_{DBD}$-DNA-Ku** | **DNA-Ku** |
| Final particle images (no.) | 436,644 | 689,183 |
| Map resolution (Å) FSC threshold 0.143 | 3.1 | 2.9 |
| Map sharpening *B* factor (Å$^2$) | 124.3 | 124.4 |
| **RealSpaceRefinement (Phenix)** | | |
| Initial model used | *MDFF refined Model* | *MDFF refined Model* |
| Model composition | | |
| Number of chains | 5 | 4 |
| Non-hydrogen atoms | 12134 | 10130 |
| Protein residues (Yku70/Yku80/$Rap1_{DBD}$) | 1386 (557/587/242) | 1144 (557/587) |
| Nucleotides | 42 (21/21) | 42 (21/21) |
| *B*-factors (Å$^2$) (min/max/mean) | | |
| Proteins | 4.51/180.22/74.00 | 0.61/168.21/64.94 |
| Nucleotides | 24.83/174.70/75.82 | 0.00/101.26/21.76 |
| R.M.S. deviations | | |
| Bond lengths (Å) | 0.003 | 0.003 |
| Bond angles (°) | 0.688 | 0.629 |
| Validation | | |
| MolProbity score | 1.86 | 2.31 |
| Clashscore | 8.09 | 7.81 |
| Ramachandran plot | | |
| Favored (%) | 93.70 | 95.88 |

| | | |
|---|---|---|
| Allowed (%) | 5.94 | 3.95 |
| Disallowed (%) | 0.36 | 0.18 |
| CCBox/CCvolume/CCmask/CCC[a] | 0.65/0.75/0.77/0.70 [a] | 0.65/0.71/0.74/0.70 [a] |
| Cβ ouliers (%) | 0.15 | 0.09 |
| CaBLAM outliers (%) | 3.20 | 2.20 |
| **Structural Comparison with starting models (RMSD CA in Å) [b]** | | |
| Yku70 structure/5Y58 chain A | 0.94 (505) / 1.16 (548) | 0.93 (485) / 1.49 (548) |
| Yku80 structure/5Y58 chain B | 0.76 (522) / 1.21 (560) | 0.97 (481) / 1.99 (560) |
| $Rap1_{DBD}$ structure/3UKG chain A | 0.94 (161) / 2.64 (223) | - |
| Yku70/Yku80 structure/5Y58 chain A/B | 0.89 (1016) / 1.25 (1108) | 1.01 (944) / 1.79(1108) |
| **Interface area in the Ku/DNA/$Rap1_{DBD}$ refined structure (Å$^2$)** | | |
| Yku70 - Yku80 | 10923 | 10232 |
| Ku - 21mer dsDNA | 1021 | 1721 |
| $Rap1_{DBD}$ - 21mer dsDNA | 2077 | - |
| $Rap1_{DBD}$ - Ku | 500 (49/455) [c] | - |

[a] Cross correlated coefficient calculated between a map simulated using the cartesian coordinates of the refined model and the experimental cryo-EM map.

[b] Superimposition achieved using the matchmaker command of *ChimeraX* 1.7.1. The RMSD and the number into parenthesis correspond to the residues selected by the matchmaker algorithm to superimpose the two structures. The second RMSD and the number of CA pairs into parenthesis correspond to the all set of CA atoms in the X-ray structures to be compared to the model ones.

[c] In parenthesis the buried area of $Rap1_{DBD}$ due to Yku70 and Yku80 respectively.

**Table S3. Yeast strains used in this study.** All strains derived from the *W303-1a* background (*ade2-1 trp1-1 ura3-1 leu2-3,112 his3-11,15 can1-100 RAD5*).

| Name | Genotype | Origin | Figure |
|---|---|---|---|
| F1092 | *MATa bar1Δ lys2::pGAL-ISCE1 ISCEI::URA3::ISCEI* | Roisne-Hamelin *et al.* 2021 | 1C |
| F1098 | *MATa bar1Δ lys2::pGAL-ISCE1 ISCEI::URA3::ISCEI rif2::HPH hml::NAT sir4::LEU2* | Roisne-Hamelin *et al.* 2021 | 1C, S4 |
| F1689 | *MATa bar1-Δ lys2::pGAL-ISCE1 IsceI::URA3::IsceI::1Rap1 site-d11::KAN* | this study | 1C, S4 |
| F1671 | *MATa bar1-Δ lys2::pGAL-ISCE1 IsceI::URA3::IsceI::1Rap1 site-d11::KAN ade2 rif2::HPH hml::NAT sir4::LEU2* | this study | 1C |
| YSM123 | *MATa bar1-Δ lys2::pGAL-ISCE1 IsceI::URA3::IsceI::1Rap1 site-d19::KAN ade2 rif2::HPH hml::NAT sir4::LEU2* | this study | 1C |
| YSM124 | *MATa bar1-Δ lys2::pGAL-ISCE1 IsceI::URA3::IsceI::1Rap1 site-d16::KAN ade2 rif2::HPH hml::NAT sir4::LEU2* | this study | 1C |
| YSM125 | *MATa bar1-Δ lys2::pGAL-ISCE1 IsceI::URA3::IsceI::1Rap1 site-d13::KAN ade2 rif2::HPH hml::NAT sir4::LEU2* | this study | 1C |
| F1674 | *MATa bar1-Δ lys2::pGAL-ISCE1 IsceI::URA3::IsceI::1Rap1 site-d23::KAN ade2 rif2::HPH hml::NAT sir4::LEU2* | this study | 1C |
| F1676 | *MATa bar1-Δ lys2::pGAL-ISCE1 IsceI::URA3::IsceI::1Rap1 site-d26::KAN ade2 rif2::HPH hml::NAT sir4::LEU2* | this study | 1C |
| F1679 | *MATa bar1-Δ lys2::pGAL-ISCE1 IsceI::URA3::IsceI::1Rap1 site-d29::KAN ade2 rif2::HPH hml::NAT sir4::LEU2* | this study | 1C |
| YSM55 | *MATa bar1-Δ lys2::pGAL-ISCE1 IsceI::URA3::IsceI::1Rap1 site-d31::KAN ade2 rif2::HPH hml::NAT sir4::LEU2* | this study | 1C |
| YSM56 | *MATa bar1-Δ lys2::pGAL-ISCE1 IsceI::URA3::IsceI::1Rap1 site-d41::KAN ade2 rif2::HPH hml::NAT sir4::LEU2* | this study | 1C |
| YSM147 | *MATa bar1-Δ lys2::pGAL-ISCE1 IsceI::URA3::IsceI::1Rap1 site- alt1-d11::KAN ade2 rif2::HPH hml::NAT sir4::LEU2* | this study | S4 |
| YSM148 | *MATa bar1-Δ lys2::pGAL-ISCE1 IsceI::URA3::IsceI::1Rap1 site-alt2-d11::KAN ade2 rif2::HPH hml::NAT sir4::LEU2* | this study | S4 |
| YSM149 | *MATa bar1-Δ lys2::pGAL-ISCE1 IsceI::URA3::IsceI::1Rap1 site-alt3-d11::KAN ade2 rif2::HPH hml::NAT sir4::LEU2* | this study | S4 |
| YSM150 | *MATa bar1-Δ lys2::pGAL-ISCE1 IsceI::URA3::IsceI::1Rap1 site-HMRE-d11::KAN ade2 rif2::HPH hml::NAT sir4::LEU2* | this study | S4 |
| YSM151 | *MATa bar1-Δ lys2::pGAL-ISCE1 IsceI::URA3::IsceI::1Rap1 site-TEF2II-d11::KAN ade2 rif2::HPH hml::NAT sir4::LEU2* | this study | S4 |

| | | | |
|---|---|---|---|
| YSM152 | *MATa bar1-Δ lys2::pGAL-ISCE1 IsceI::URA3::IsceI::1Rap1 site-RPS17-d11::KAN ade2 rif2::HPH hml::NAT sir4::LEU2* | this study | 4S |
| YSM153 | *MATa bar1-Δ lys2::pGAL-ISCE1 IsceI::URA3::IsceI::1Rap1 site-RPS11B-d11::KAN ade2 rif2::HPH hml::NAT sir4::LEU2* | this study | 4S |
| YSM154 | *MATa bar1-Δ lys2::pGAL-ISCE1 IsceI::URA3::IsceI::1Rap1 site-RPL41II-d11::KAN ade2 rif2::HPH hml::NAT sir4::LEU2* | this study | 4S |
| F1705 | *MATa bar1-Δ lys2::pGAL-ISCE1 IsceI::URA3::IsceI::1Rap1 site-mutated1-d11::KAN ade2 rif2::HPH hml::NAT sir4::LEU2* | this study | 1C |
| F1707 | *MATa bar1-Δ lys2::pGAL-ISCE1 IsceI::URA3::IsceI::1Rap1 site-mutated2-d11::KAN ade2 rif2::HPH hml::NAT sir4::LEU3* | this study | 1C |
| F1713 | *MATa bar1-Δ lys2::pGAL-ISCE1 IsceI::URA3::IsceI::1Rap1 site-mutated3-d11::KAN ade2 rif2::HPH hml::NAT sir4::LEU4* | this study | 1C |
| YSM204 | *MATa bar1-Δ lys2::pGAL-ISCE1 IsceI::URA3::IsceI::1Rap1 site-d11 ade2 rif2::HPH hml::NAT sir4::LEU2 KU70-13MYC:KAN* | this study | 1D, S6 |
| YSM205 | *MATa bar1-Δ lys2::pGAL-ISCE1 IsceI::URA3::IsceI::1Rap1 site-mutated3-d11 ade2 rif2::HPH hml::NAT sir4::LEU2 KU70-13MYC:KAN* | this study | 1D, S6 |
| Lev1802 | *MATa bar1Δ lys2::pGAL-ISCE1 ISCEI::URA3::ISCEI yku80::TRP1* | this study | S20B |
| Lev1804 | *MATa bar1-Δ lys2::pGAL-ISCE1 IsceI::URA3::IsceI::1Rap1 site-d11::KAN ade2 rif2::HPH hml::NAT sir4::LEU2 yku80::TRP1* | this study | S20B |
| Lev1559 | *MATa proLEU2-loxP-CEN6-klURA3-loxP-orfLEU2-pGAL1-CRE::NAT* | Pobiega *et al.* 2021 | 4C, S2B-C S21B-D |
| Lev1868 | *MATa proLEU2-loxP-CEN6-klURA3-loxP-orfLEU2-pGAL1-CRE::NAT lig4::HIS3* | this study | S2B-C |
| Lev1326 | *MATa loxP-NAT-CEN6-klURA3-loxP lys2::pGAL1-CRE::NAT rif2::HPH sir4::HPH hml::NAT* | Pobiega *et al.* 2021 | 4B-C S21B-D |
| Lev1181 | *MATa loxP-klTRP1-CEN6-loxP-skHIS3 rap1-(Δ)::KAN rif2::HPH sir4::HPH hml::NAT* | this study | 4B-C S21B-D |
| Lev608 | *MATa bar1 tel1::HIS3 rif2::HPH sir4::HPH hml::NAT* | Marcand *et al.* 2008 | S22A-B |
| Lev707 | *MATa bar1 tel1::HIS3 rif2::HPH sir4::HPH hml::NAT rap1-(Δ)::KAN* | Marcand *et al.* 2008 | S22A-B |
| Lev1810 | *MATa bar1 tel1::HIS3 rif2::HPH sir4::HPH hml::NAT pol4::TRP1* | this study | S22A-B |

### SUPPLEMENTARY FIGURES

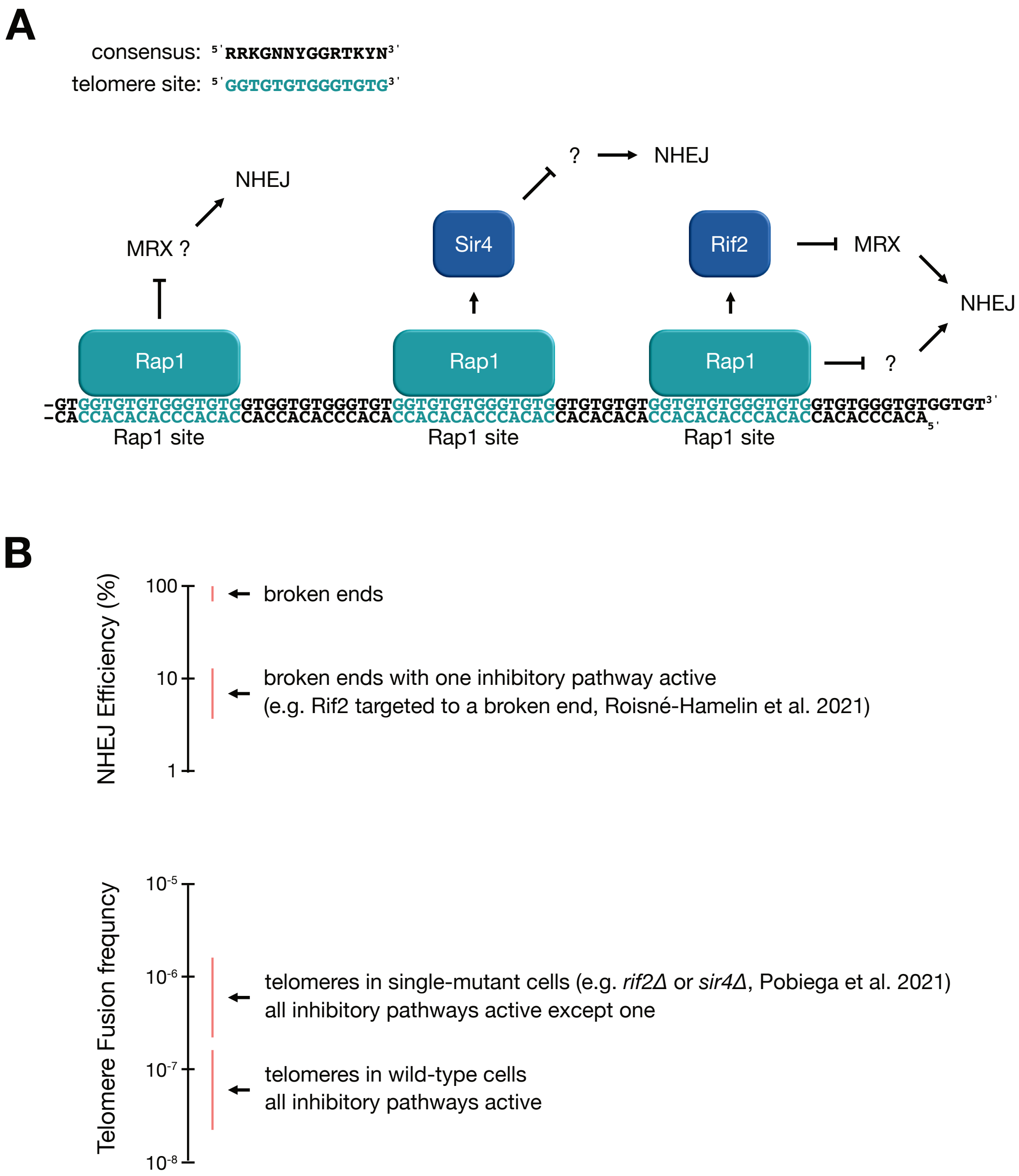


**Supplementary Figure S1. Telomere end protection against fusion in *S. cerevisiae*.** (**A**) (top) Rap1 binding site consensus shown in black (R: G or A, K: G or T, Y: T or C, N: any base) (Buchman et al. 1988) and Rap1 telomere site in teal. (bottom) Schematic representation of the pathways blocking NHEJ at telomeres. Rap1 binds to the double-stranded telomere repeats in tandem and recruits several co-factors to telomeres, including Rif2 and Sir4 which inhibits NHEJ. Rif2 exerts its effect by directly inhibiting the recruitment of the Mre11-Rad50-Xrs2 complex (MRX) (Roisné-Hamelin et al. 2021, Khayat et al. 2021). Sir4 inhibits NHEJ at telomeres through an unknown mechanism (Pobiega et al. 2021, Bordelet et al. 2023). Independently of Rif2 and Sir4, the binding of multiple Rap1 in tandem may also block MRX recruitment, and consequently, may inhibit NHEJ (Negrini et al. 2007, Marcand et al. 2008, Lescasse et al. 2013). In the present study, we address the contribution of the end-proximal Rap1 to telomere protection against NHEJ. (**B**) Schema illustrating the relative contribution of individual pathways to NHEJ inhibition at telomeres. The loss of one pathway has a relatively mild effect, increasing the fusion frequency only a few-fold above the basal frequency observed in wild-type cells (e.g., approximately 3-fold in *rif2Δ* cells, using the CFC assay (Pobiega et al. 2021), see also **Fig. S2** for a schema of the assay). Conversely, tethering Rif2 alone at a broken end only partially inhibits NHEJ repair (approximately an order of magnitude less efficient in an IScel assay (Roisné-Hamelin et al. 2021), same assay than in **Fig. 1B-C**).

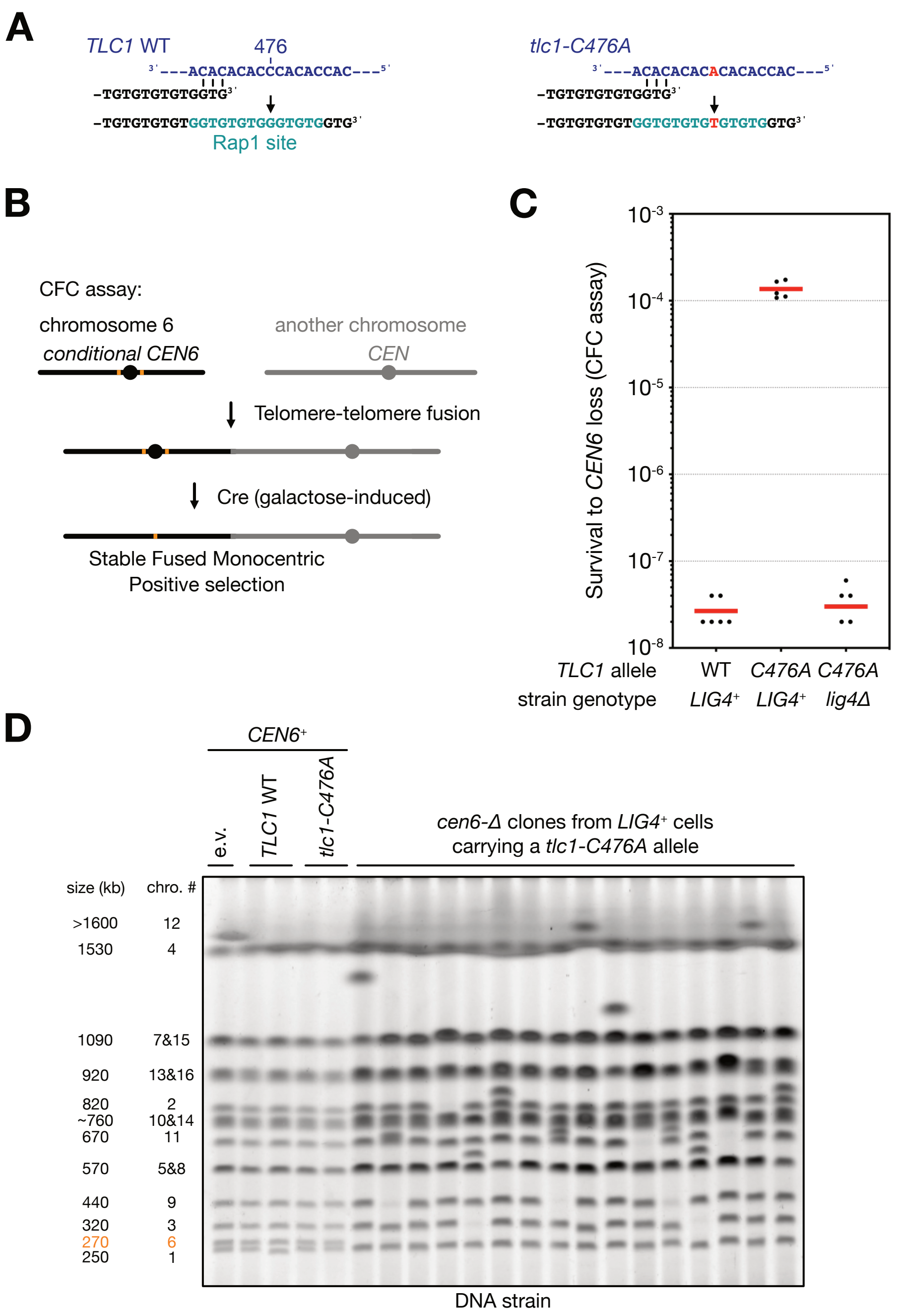


**Supplementary Figure S2. Frequent NHEJ-dependent chromosome end fusions in wild-type cells carrying an ectopic copy of the *tlc1-C476A* allele.** (**A**) Wild-type and *C476A* mutant template sequences of the *S. cerevisiae* telomerase RNA *TLC1*. Copying the *C476A* template generates telomere repeats devoid of the 5'-TGGGT-3' core motif, which is essential for Rap1 high-affinity binding (Prescott and Blackburn 2000, Chan et al. 2001). (**B**) Schema of the Chromosome Fusion Capture assay (CFC) (Pobiega et al. 2021). In this assay, the relative survival to the loss of a native centromere (*CEN6*) serves as a proxy for chromosome end fusion frequency. (**C**) Survival to *CEN6* loss in wild-type (*LIG4+*) and NHEJ-deficient (*lig4Δ*) cells carrying an extra-copy of the wild-type (WT) or mutant (*C476A*) *TLC1* gene on a centromeric plasmid (pRS314). The presence of the mutant *TLC1-C476A* allele increases the frequency of NHEJ-dependent fusion events by more than 1000-fold. (**D**) Karyotype of *cen6Δ* clones from cells carrying a *tlc1-C476A* allele. In each clone, chromosome 6 is fused to another chromosome. Overall, these results show that limiting Rap1 binding at telomere ends leads to frequent fusions in *S. cerevisiae*, as previously observed in the budding yeast *Kluyveromyces lactis* (McEachern et al. 2000), the expected consequence of a partial loss of all the inhibitory pathways established by Rap1 at telomeres (**Fig. S1**).

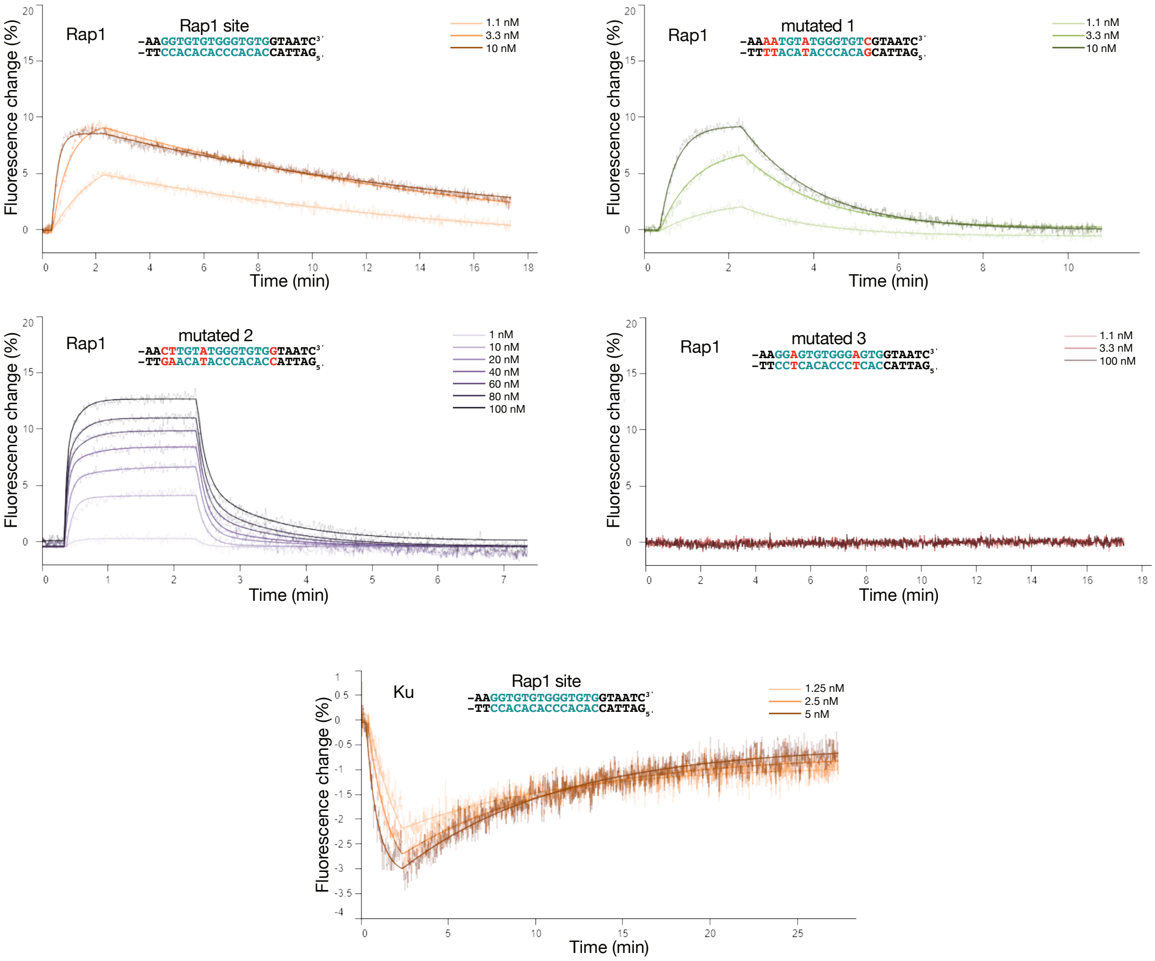


| Protein | DNA | $k_{on1}$ ($M^{-1}s^{-1}$) | $k_{on2}$ ($M^{-1}s^{-1}$) | $k_{off1}$ ($s^{-1}$) | $k_{off2}$ ($s^{-1}$) | $k_{d1,1}$ (nM) | $k_{d2,2}$ (nM) |
|---|---|---|---|---|---|---|---|
| Rap1 | Rap1 site | 8.2 ± 2.0 E+6 | - | 1.1 ± 0.2 E-3 | - | 0.15 ± 0.06 | - |
| Rap1 | mutated 1 | 2.2 ± 0.9 E+6 | - | 8.6 ± 0.7 E-3 | - | 4.4 ± 1.2 | - |
| Rap1 | mutated 2 | 3.9 ± 1.2 E+6 | 6.8 ± 1.9 E+5 | 1.3 ± 0.1 E-1 | 1.2 ± 0.3 E-2 | 38 ± 13 | 19 ± 3 |
| Rap1 | mutated 3 | N/A | - | N/A | - | N/A | - |
| Ku | Rap1 site | 5.3 ± 1.1 E+6 | - | 2.1 ± 0.6 E-3 | - | 0.39 ± 0.20 | - |

**Supplementary Figure S3. Rap1-DNA and Ku-DNA affinities measure by switchSENSE**. Rap1 site and mutated sequences were hybridized to switchSENSE DNA-adapters as dsDNA overhangs ("ligand"). A fluorescent dye on the adapter serves as a reporter fluorophore to detect binding events of proteins ("analytes") to these ligands causing the signal to increase or decrease. The sensorgram for each interaction shows the association and dissociation of Rap1 or Ku with the respective dsDNA sequence. The kinetic parameters (on-rate, off-rate) were obtained by applying a global fit to curves measured at multiple concentrations. For Rap1 binding to the native Rap1 site, a triple digit picomolar affinity was measured, while for the mutated sequence 1, a lower affinity of around 5 nM was determined. The interaction of Rap1 with the mutated sequence 2 yielded a biphasic interaction profile indicating two different or consecutive binding events taking place. The affinity for these interactions were recorded to be in the double-digit nanomolar range. No binding was observed for Rap1 to the mutated sequence 3. For Ku binding to the Rap1 site, a low triple-digit picomolar affinity was measured. All experiments were carried out in triplicate and the results averaged.

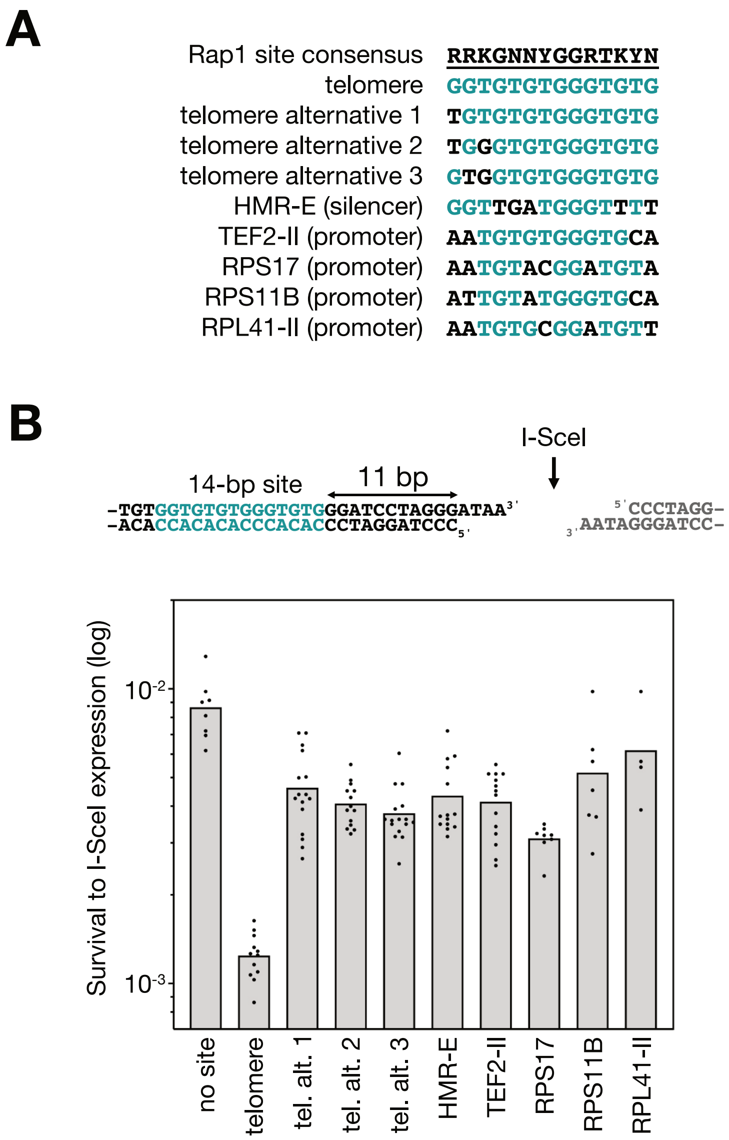


**Supplementary Figure S4. NHEJ inhibition by native Rap1 binding sites.** (**A**) Sequences of the sites tested in this study. The Rap1 binding site consensus (Buchman et al. 1988) is underlined (R: G or A, K: G or T, Y: T or C, N: any base). Bases identical to the telomere site sequence in teal. Bases distinct from the telomere site in black. The telomere site matching the consensus is identical to the one used in Fig. 1. The three alternative telomere sites are present within native telomere sequences and originate from the alternative priming positions of yeast telomerase (see **Fig. S5**). Other tested native sites are derived from a silencer element (*HMR-E*) and from gene promoters (*TEF2*, *RPS17*, *RPS11* and *RPL14*) which are bound by Rap1 *in vivo* (Buchman et al. 1988, Challal et al. 2018). (**B**) Native Rap1 sites diverging from the telomere site are less effective at blocking NHEJ at a broken end. 11 bp separate the edge of the sites from the broken end. Cells lacking Rif2 and Sir4. Mean from independent cell cultures. Statistical analysis in **Table S1**.

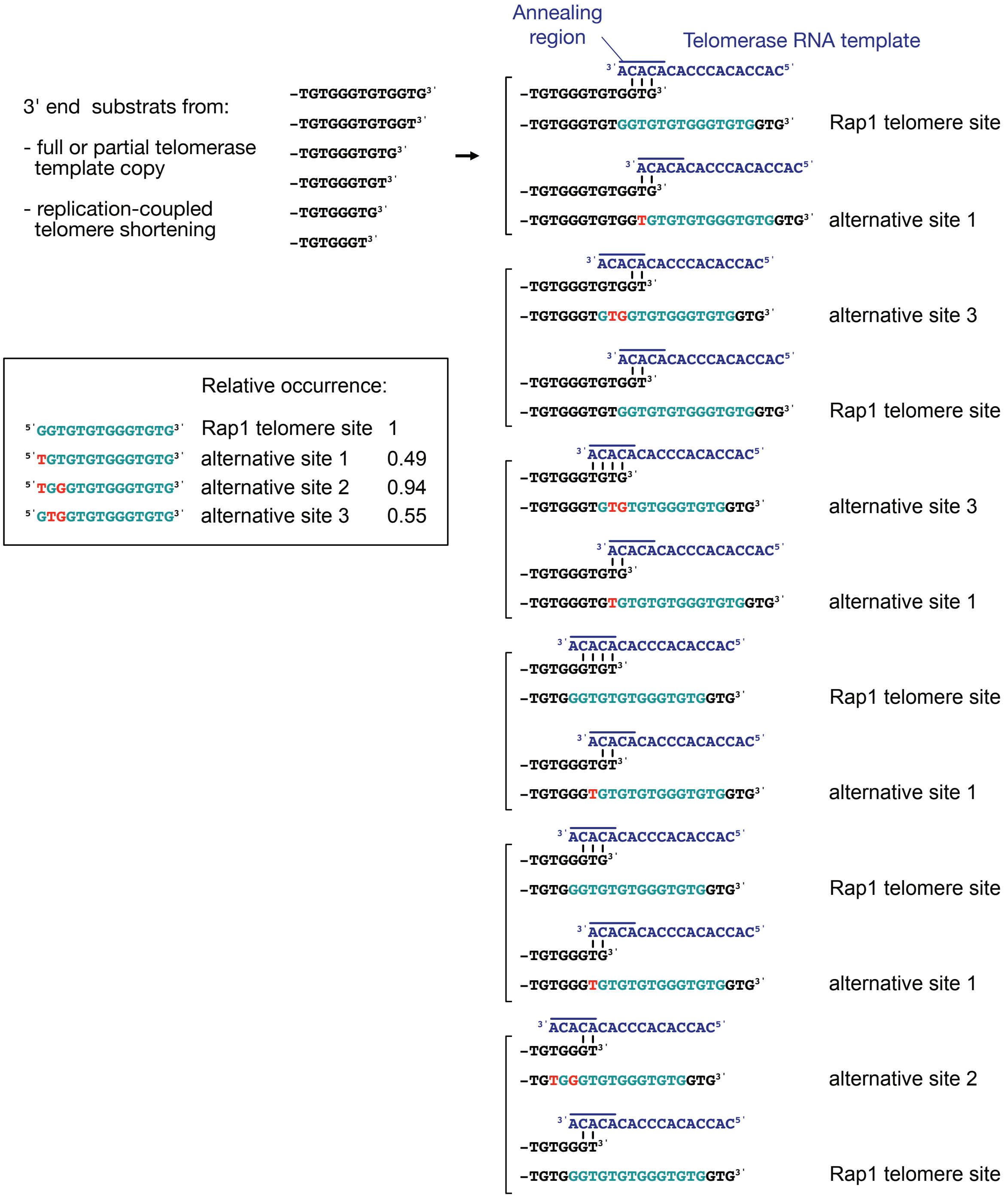


**Supplementary Figure S5. Products of *S. cerevisiae* telomerase DNA synthesis.** Multiple priming positions on the yeast telomerase RNA template generate telomere sequence variability (Förstemann and Lingner 2001). Telomerase DNA synthesis produces Rap1 telomere sites matching the consensus sequence, as well as alternative sites with one or two mismatches. The relative occurrence of the alternative sites was determined using sequences of unfused telomeres from wild-type cells (**Fig. 4C**). Bases identical to the telomere site sequence in teal. Bases distinct from the telomere site in red.

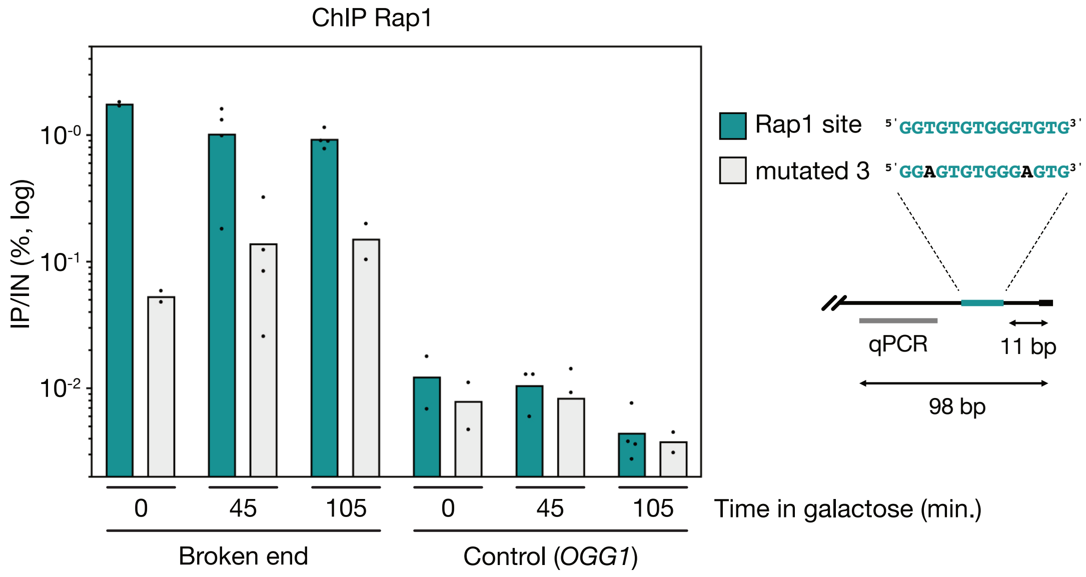


**Supplementary Figure S6. ChIP analysis of Rap1 binding at broken ends.** Rap1 binding at a broken end with a Rap1 site (teal) or a mutated site (light gray) determined by ChIP. Cells were arrested in G1 prior to induction of the I-SceI endonuclease by galactose addition to the medium. Rap1 site 11 bp from the broken end. Quantification of immunoprecipitated DNA (IP) relative to the input DNA (IN). Means from independent samples. Control within *OGG1* coding sequence. The interaction of Rap1 with the mutated site is reduced compared to its interaction with the Rap1 telomere site.

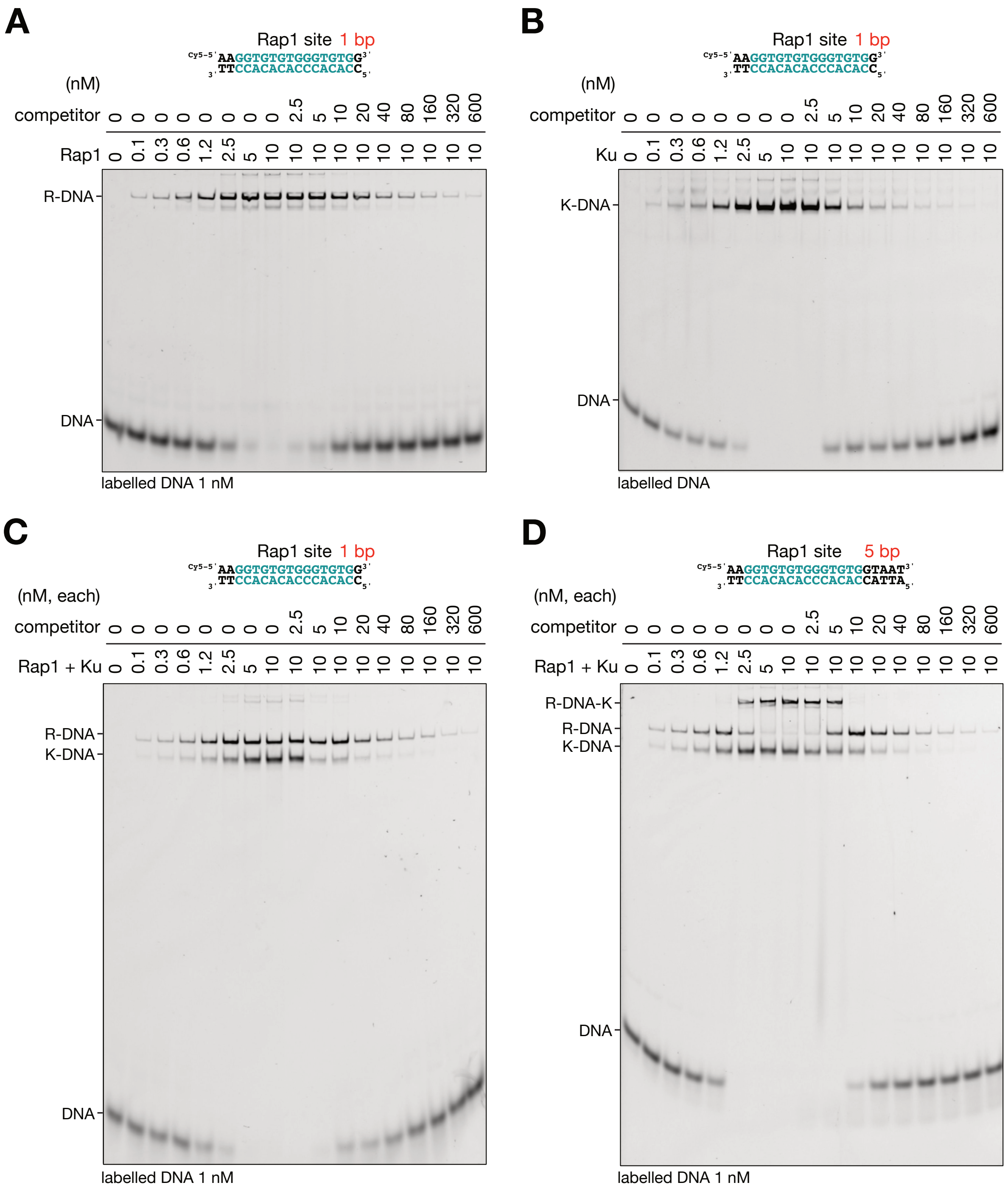


**Supplementary Figure S7.** EMSA analysis of Rap1 and Ku binding on duplex DNA. (**A-D**) Representative EMSA results of Rap1 and Ku binding on Cy5-labelled DNA duplexes with a single Rap1 site and a 1 or 5 bp double-strand DNA extension downstream of the site (labelled DNA at a fixed concentration of 1 nM). Unlabelled competitor: duplex DNA with a Rap1 site and a downstream double-stranded DNA extension of 5 bp.

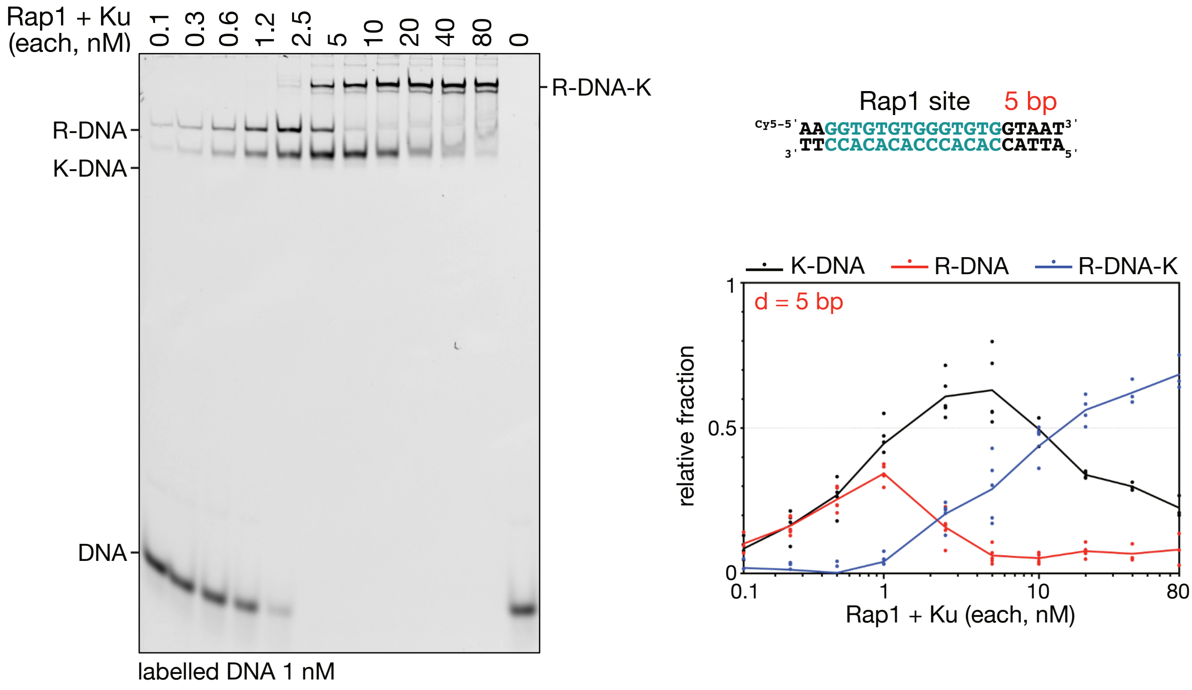


**Supplementary Figure S8.** EMSA analysis of Rap1 and Ku binding on duplex DNA. *Left,* representative titration of Rap1 and Ku binding to a DNA duplex with a 5 bp extension downstream of the Rap1 site (labelled DNA at a fixed concentration of 1 nM). *Right,* Quantified EMSA data showing the association of Rap1 and Ku. Means from independent experiments.

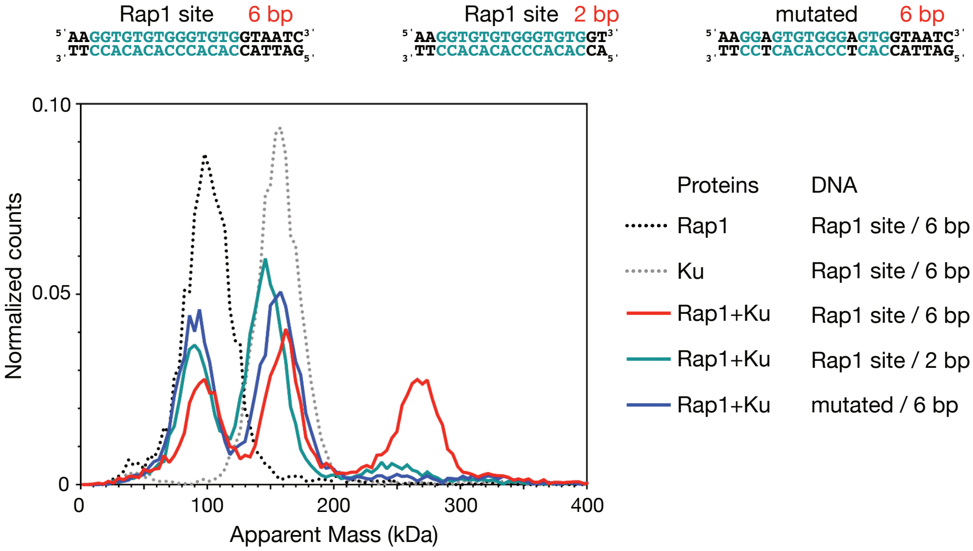


**Supplementary Figure S9. Mass photometry analysis of the Rap1-DNA-Ku ternary complex.** DNA 50 nM, proteins 50 nM. Expected molecular weights (kDa): Rap1 94, Rap1-DNA 108, Ku 144, DNA-Ku 158, Rap1-DNA-Ku 252. Complexes of the expected size for the Rap1-Ku-DNA complex assembled on DNA duplexes with a Rap1 site and 6 bp downstream extension (red) but not on duplexes with a 2 bp extension (teal) or with a mutated Rap1 site (blue). Signals from free Rap1 and Rap1-DNA complexes, and from free Ku and DNA-Ku complexes partially overlap.

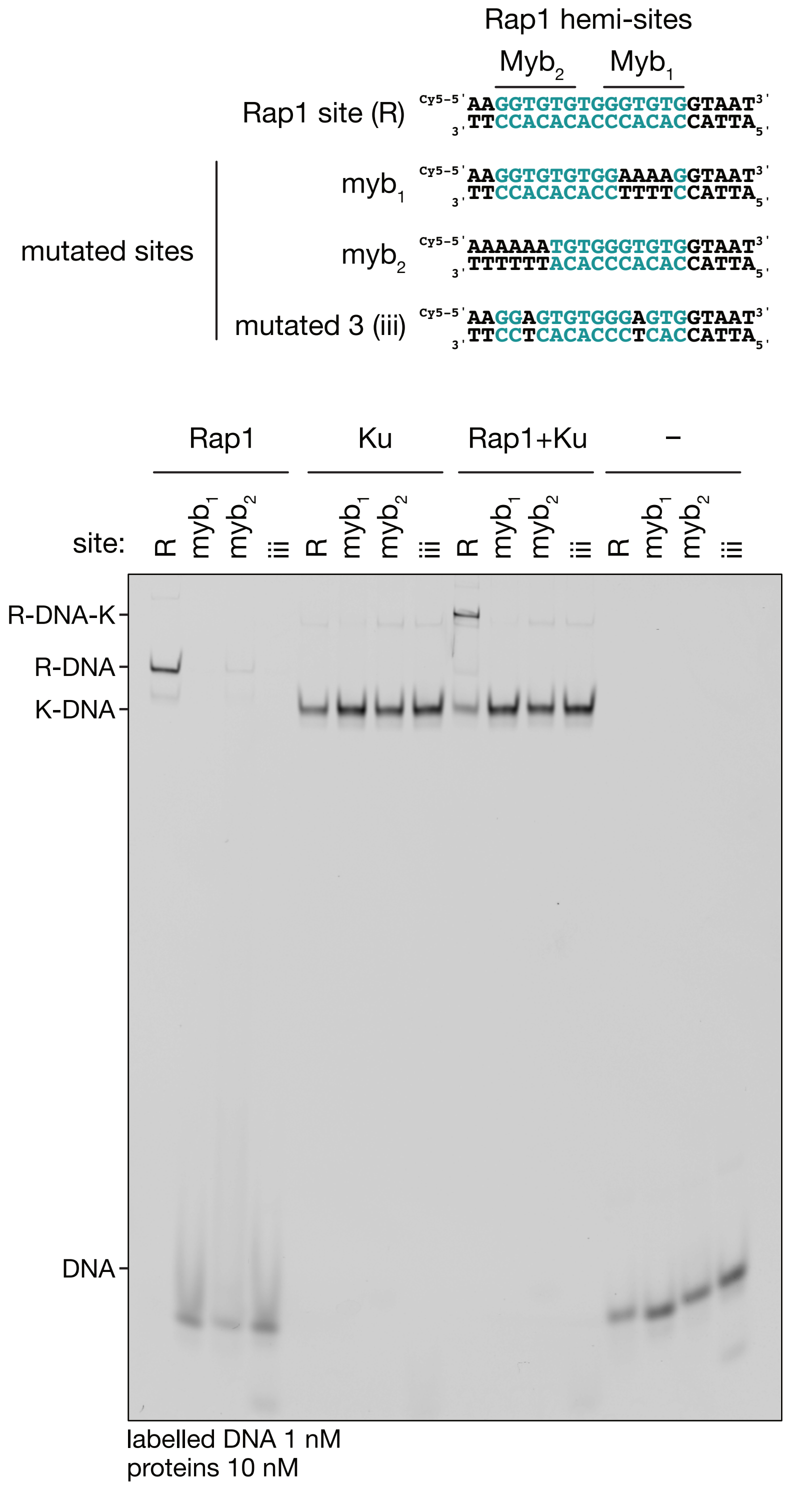


**Supplementary Figure S10. Mutations in either Rap1 hemi-site impaired both Rap1's interaction with DNA and the formation of Rap1-DNA-Ku complexes.** DNA duplexes with a 5 bp DNA extension downstream of a Rap1 or mutated site. As expected, Ku binding appears sequence-independent.

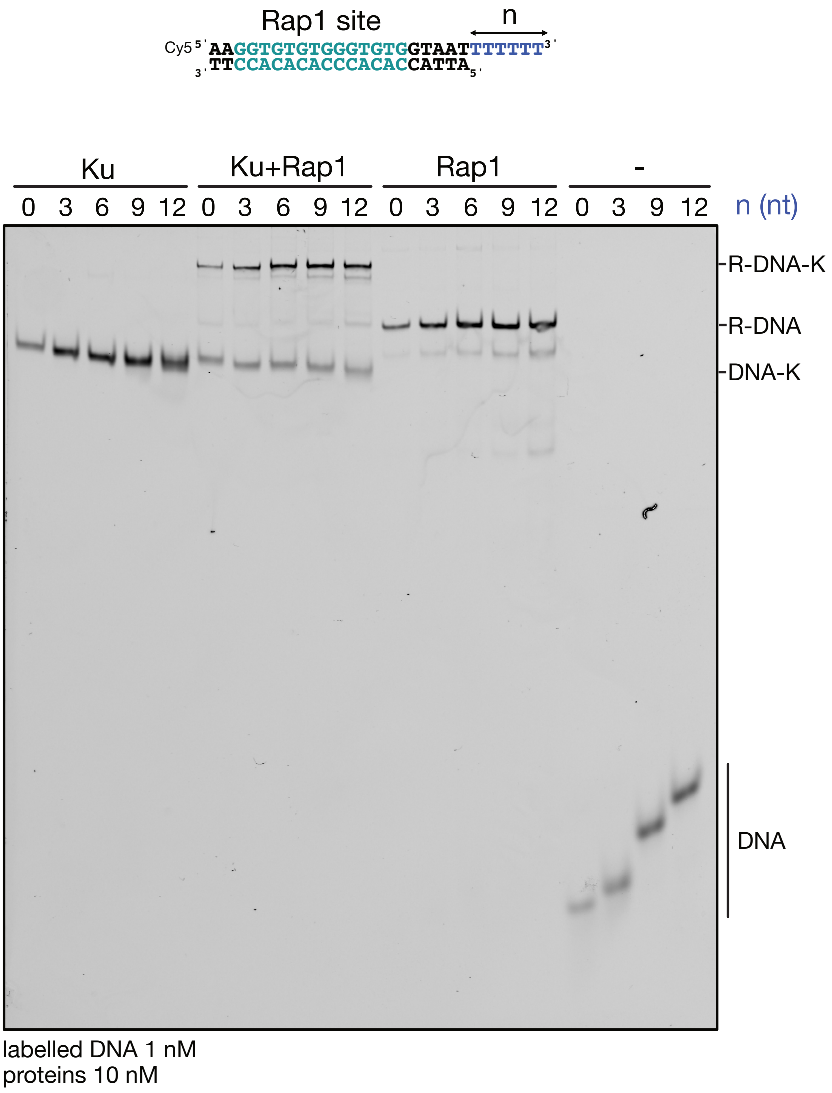


**Supplementary Figure S11. 3' single-strand overhangs do not prevent the formation of Rap1-DNA-Ku ternary complexes.** DNA duplexes with a 5 bp double-strand DNA extension downstream of the Rap1 site. *(n)* number of nucleotides of 3' single-strand overhangs (0, 3, 6, 9 and 12 nt). Ku binds DNA and Rap1-DNA complexes with an 3' overhang of up to 12 nt.

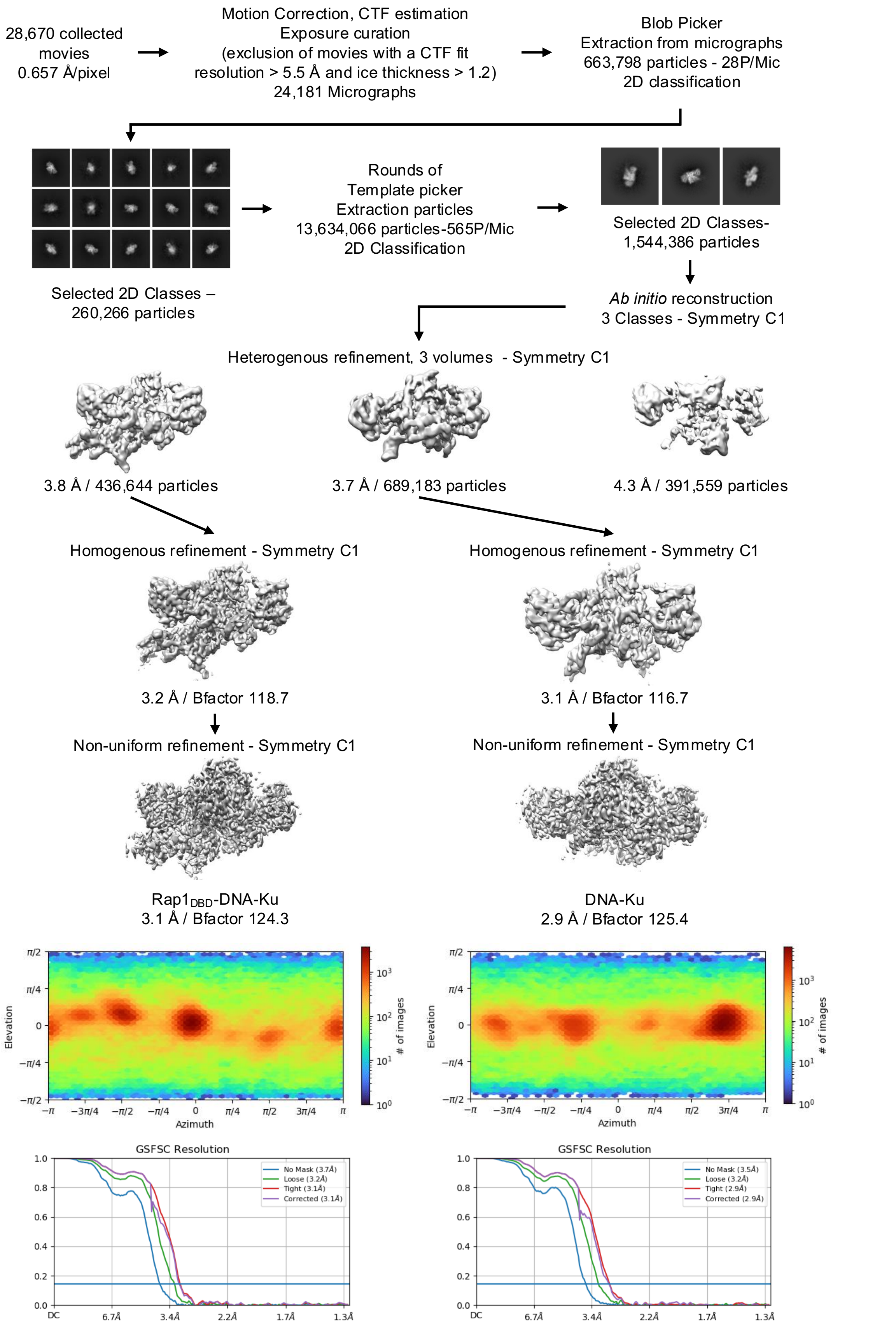


**Supplementary Figure S12. Cryo-EM data processing scheme of the ternary Rap1$_{DBD}$-DNA-Ku complex and the binary DNA-Ku complex.** The gold standard Fourier shell correlation (GSFSC) curves and orientation diagnosis plots (cryoSPARC) are given for two structures.

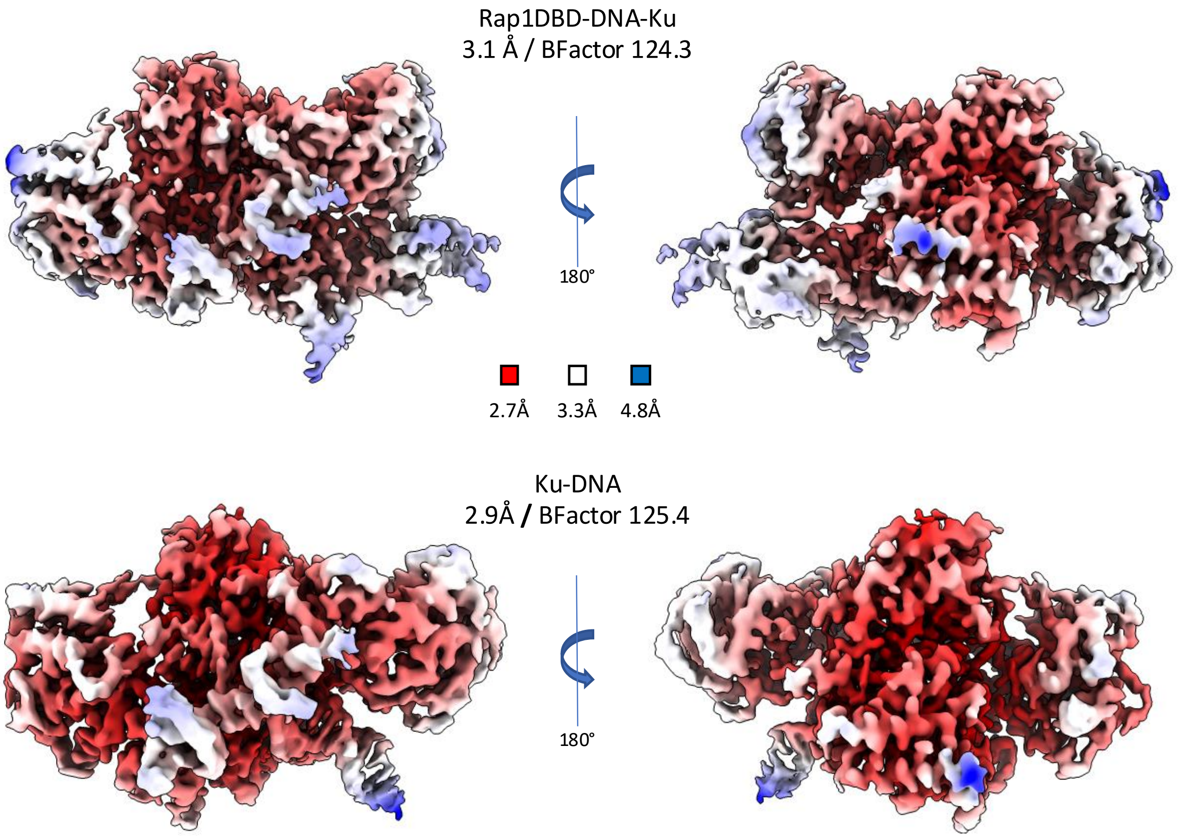


**Supplementary Figure S13**. **Local resolution maps for the Ku-DNA-Rap1 complex (top) and the Ku-DNA complex (bottom)**. Maps were calculated with the local resolution tool of Phenix. The two half maps obtained for each complex were first masked with the MapBox function of Phenix based on the respective models, to generate the masked maps. The color code is the same for the two local resolution maps.

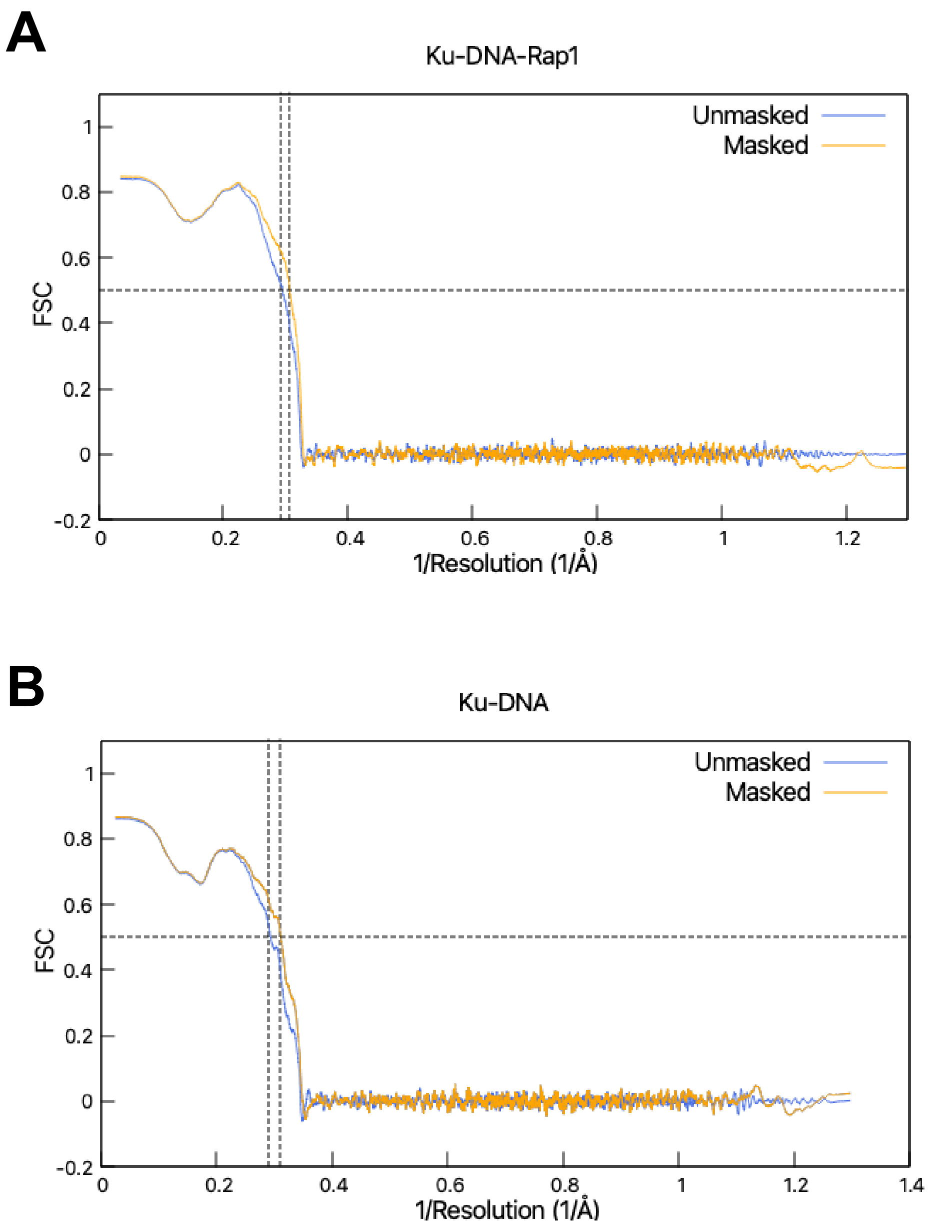


**Supplementary Figure S14. Model to Map Fourier Shell Correlations for the Ku-DNA-Rap1 model (A) and the Ku-DNA model (B).** Correlations were calculated with the validation tools of Phenix. The FSC were calculated with the unmask data (in blue) and the masked data (in orange).

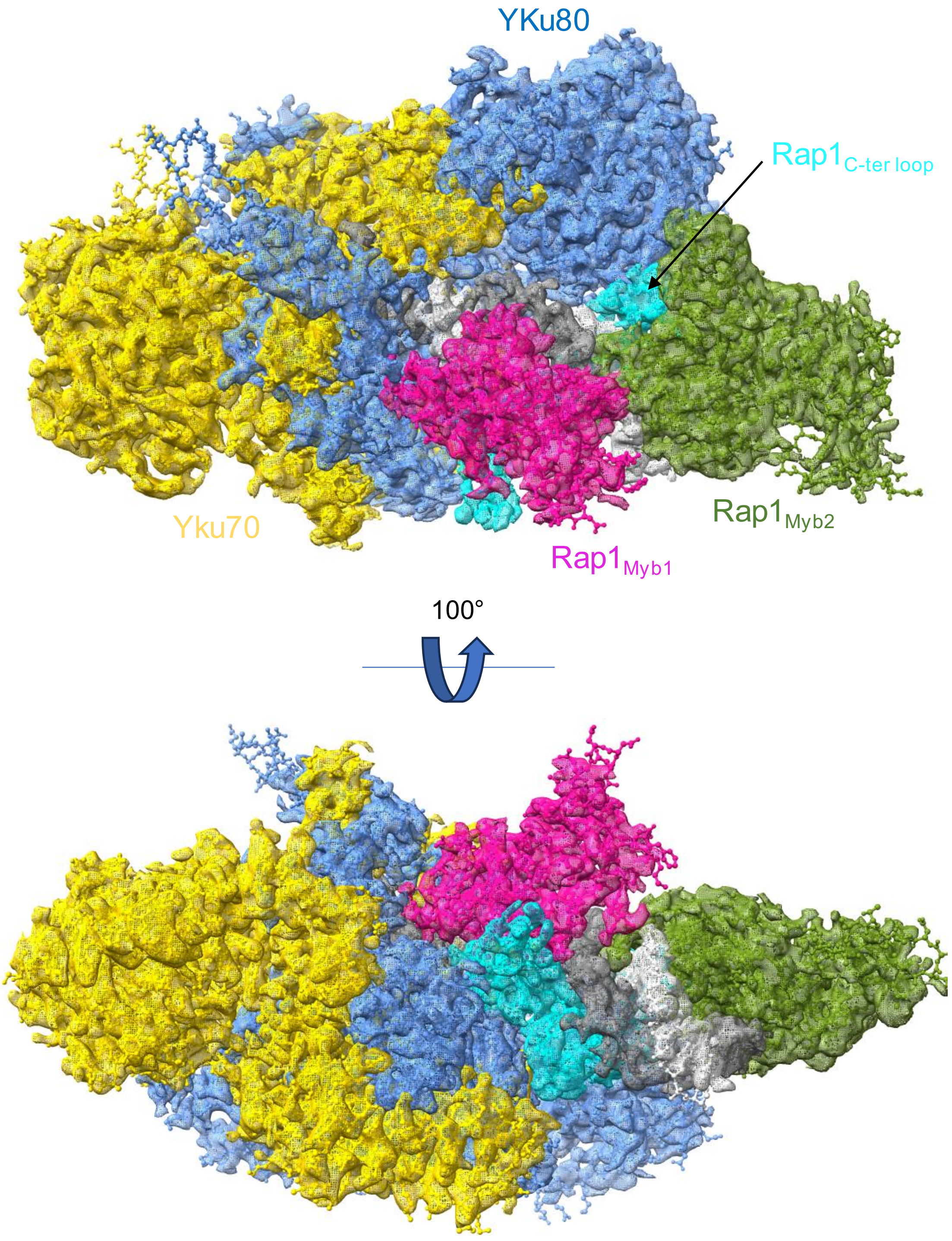


**Supplementary Figure S15. Structural agreement between the refined model and the cryo-EM map of the Rap1-DNA-Ku ternary complex.** The refined model is depicted in cartoon representation with Yku70, Yku80, the DNA G-rich strand and the DNA C-rich strand coloured in gold, cornflower blue, light grey and grey, respectively. The Rap1 subdomains are highlighted as follows: $Rap1_{Myb1}$ in pink, $Rap1_{Myb2}$ in green and $Rap1_{Cter\text{-}loop}$ in turquoise. The same colour scheme is applied to the cryo-EM map, which is shown in a semi-transparent view for comparison. At the selected threshold, electron density is absent in regions corresponding to residues 199-213 of Yku70, 166-169 and 288-301 of Yku80 and 481-504 of Rap1.

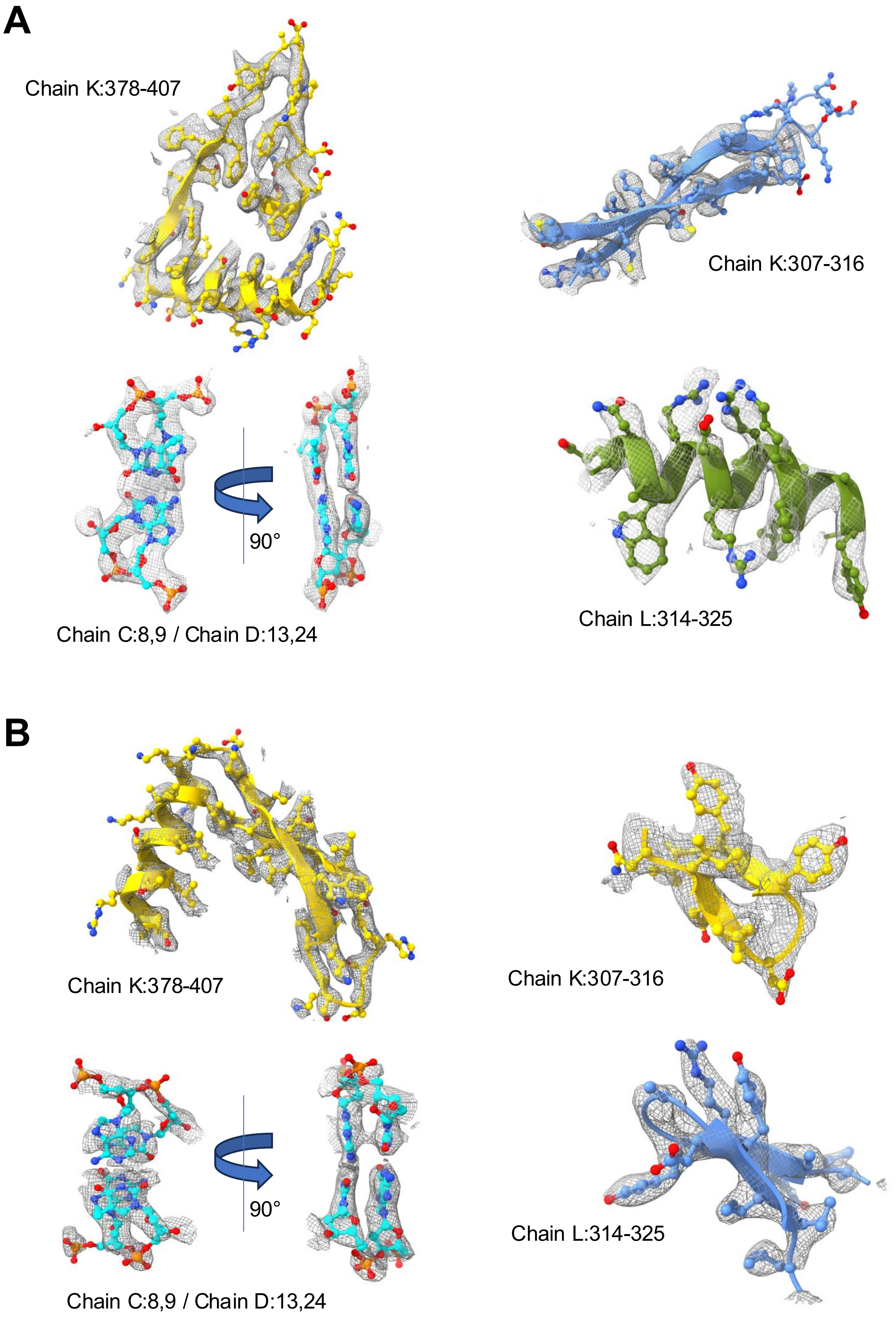


**Supplementary Figure S16. Examples of atomic models in the cryoEM maps for Ku-DNA-Rap1 complex (A) and the Ku-DNA complex (B).** The identity of the residues or nucleotides are indicated for each the segment.

A

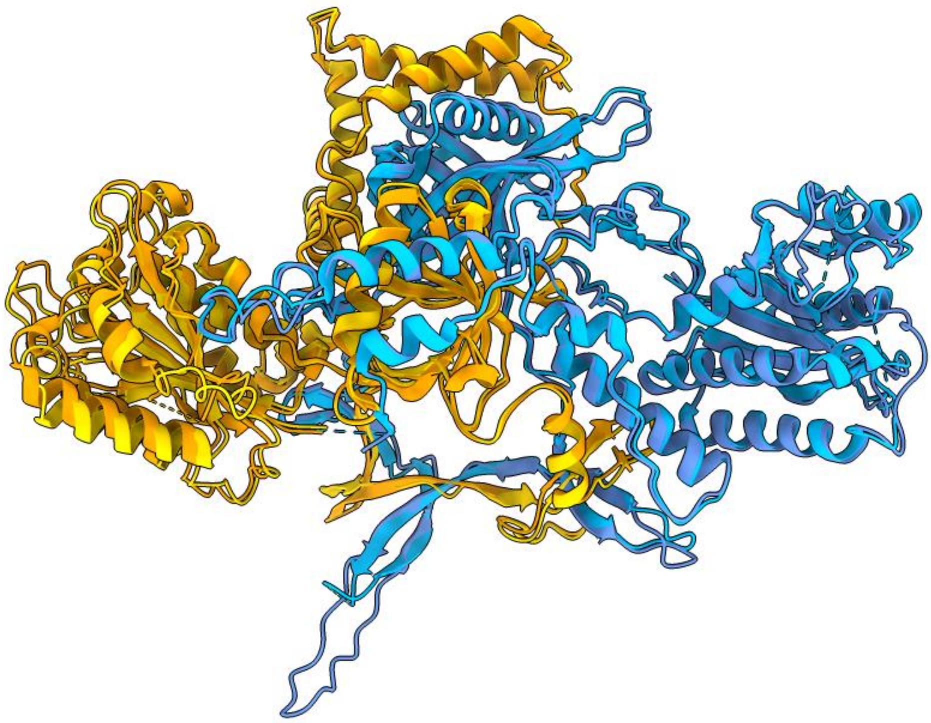

B

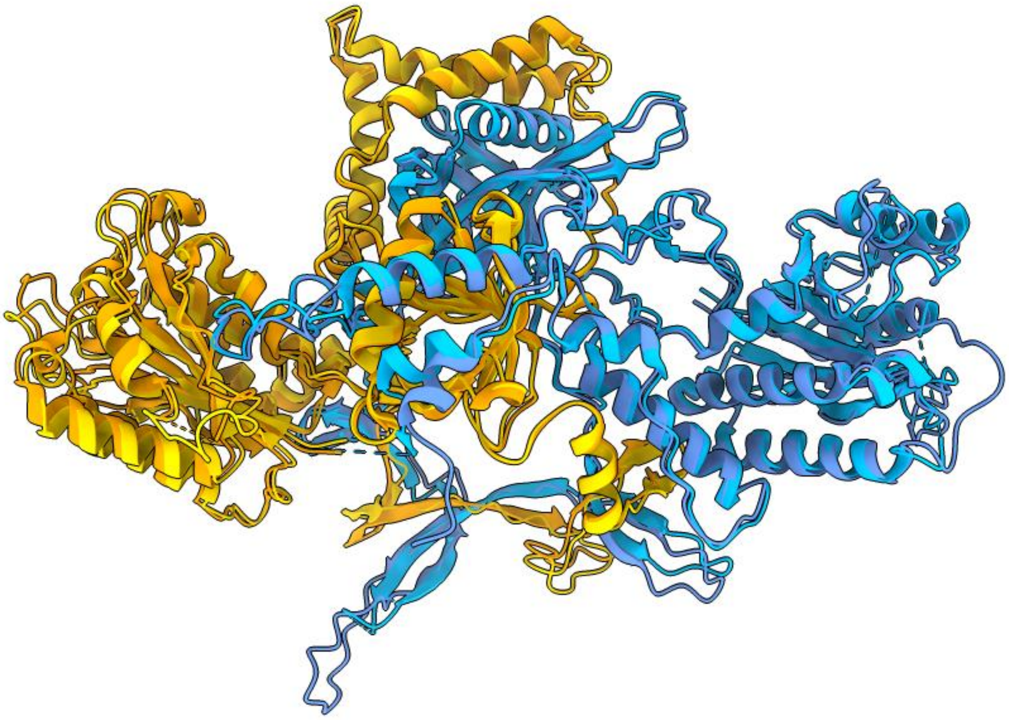

**Supplementary Figure S17. Comparison of the Ku structures in the Ku-DNA-Rap1 complex (A) and the Ku-DNA complex (B) with the yeast Ku X-ray structure 5Y58.** In (A) and (B), Yku70 and Yku80 are shown in orange and light blue respectively for the 5Y58 structure and in gold and cornflower blue for the Yku70 and Yku80 structures in the Ku-DNA-Rap1 complex and the Ku-DNA complex. The corresponding RMSD on the Ca atoms are indicated in **Table S2.**

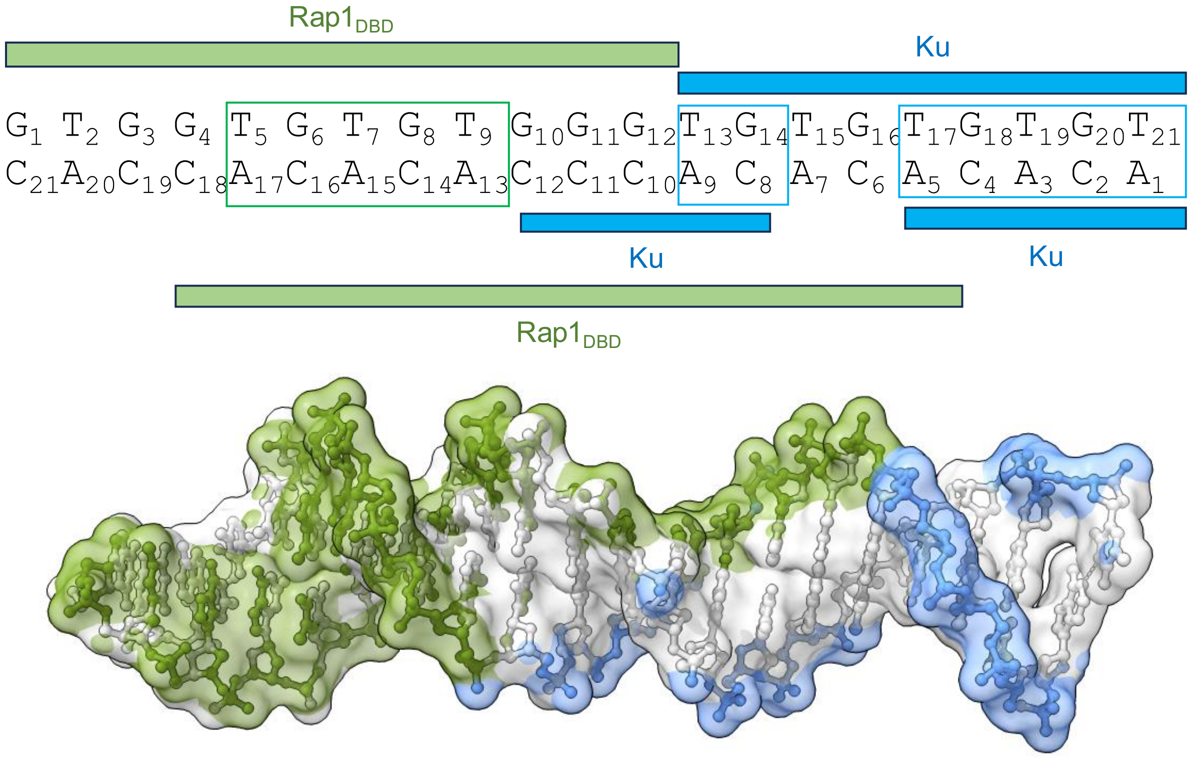


**Supplementary Figure S18. Asymmetry between the DNA binding sites of Rap1 and Ku in the Rap1-DNA-Ku ternary complex.** The DNA engaged into the Ku beta-bridge (handle) involves nucleotides T17 to T21 on the G-rich strand and the complementary nucleotides A1 to A5 on the C-rich strand. Other DNA interactions with Ku involving only one of the two strands, specifically nucleotides T13 to G16 of the G-rich strand and C8 to C12 on the C-rich strand. Rap1 interacts with a double-stranded region (G4-G12 of the G-rich strand and the complementary nucleotides C10-C18 on the C rich strand) as well as with nucleotides C6 and A7 of the C-rich strand (paired with T15 and G16 on the G-rich strand that interact with Ku) and nucleotides G11-G13 (paired with C10-C12 that interact with Ku)

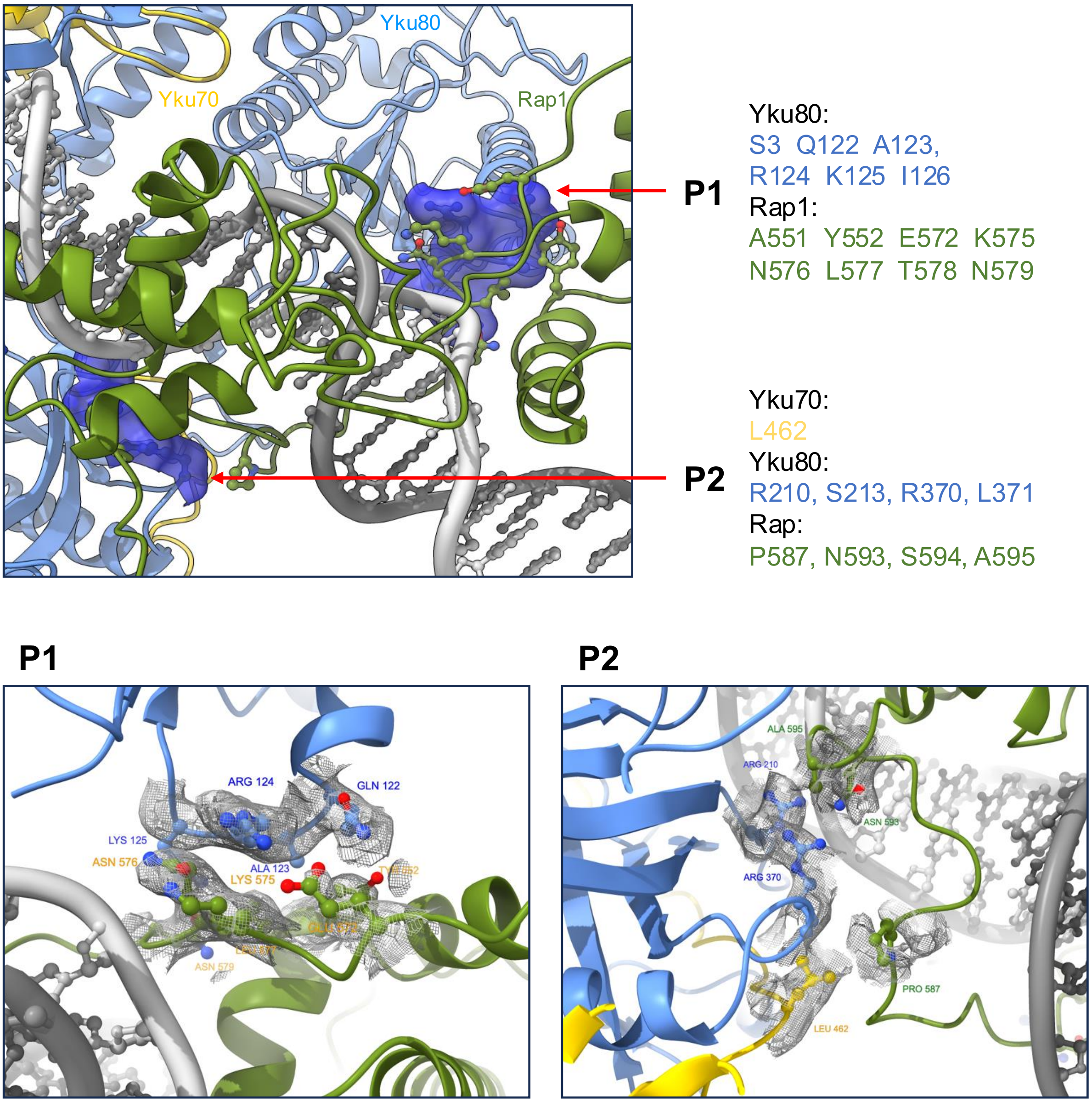


**Supplementary Figure S19. Rap1-Ku interface in the Rap1-DNA-Ku ternary complex.** The interface is restricted to two patches, P1 and P2, where the distance between the two proteins is below 4.0 Å. P1 (280 Å$^2$) involves five residues of the Myb2 subdomain of Rap1$_{DBD}$ (A551, Y552, E572, K575-N579) and six residues of Yku80 (S3 and the Q122-I126 loop). P2 (120 Å$^2$) involves four residues located in the Rap1$_{DBD}$ C-terminal wrapping loop (P587, N593-A595) and five residues Yku80 and Yku70 (L462 of yKu70 and R210, S213, R370, L371 of yKu80). Rap1$_{DBD}$, Yku70, Yku80, DNA G-rich strand and the DNA C-rich strand are show in cartoon representation and coloured in green, gold, cornflower blue, light grey and grey, respectively. The bottom insets display the local map of the Rap1-Ku interface with the sidechains of the patches shown in ball-and-stick representation.

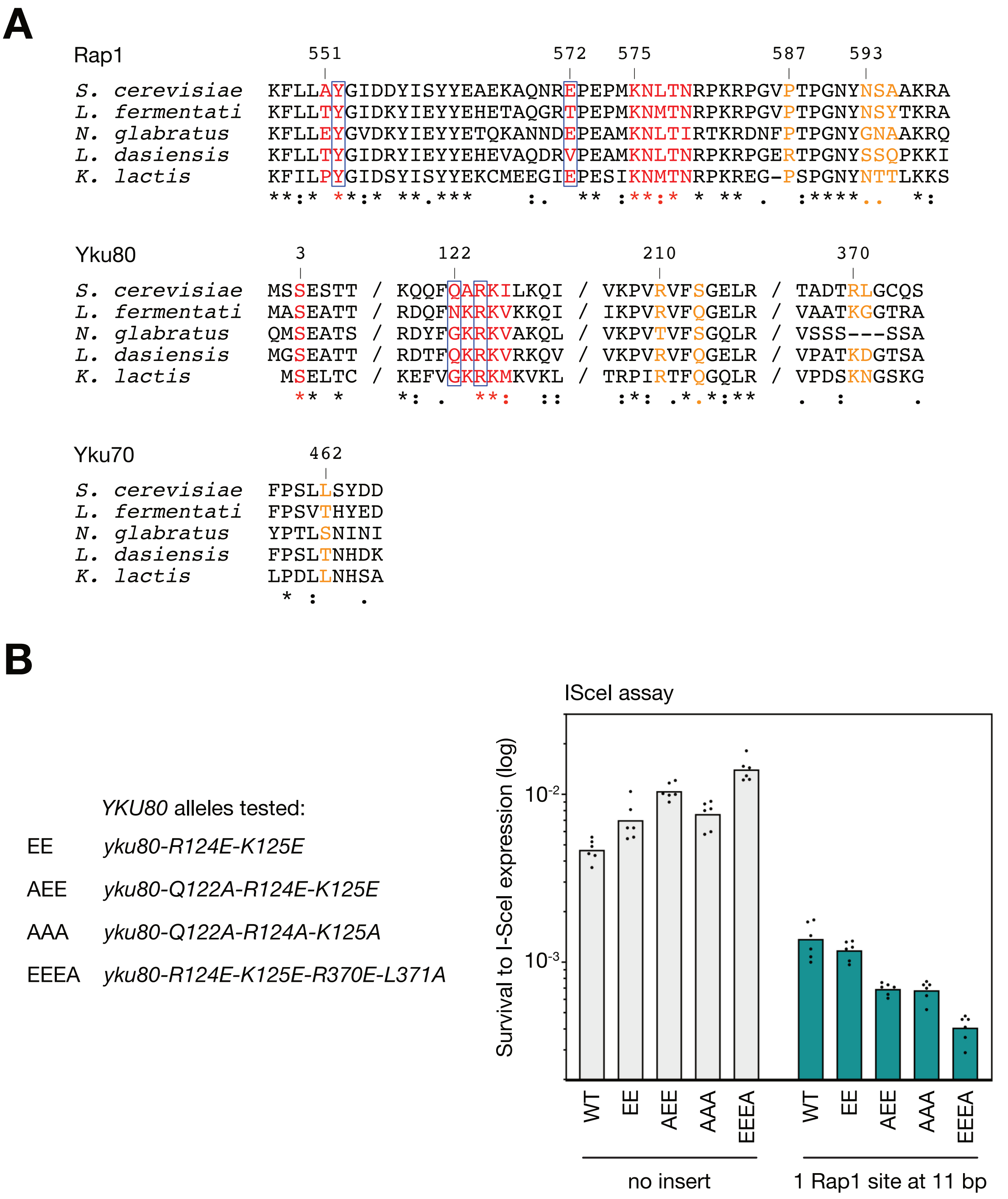


**Supplementary Figure S20. Conservation and mutagenesis of the Rap1-Ku interface in the Rap1-DNA-Ku ternary complex.** (**A**) Multiple sequence alignments of Rap1, Yku80 and Yku70 across yeast species. The indicated positions correspond to the sequences from *S. cerevisiae*. Residues in patch 1 and patch 2 are highlighted in red and orange, respectively. Residues that could contribute to hydrogen bond or salt bridge intermolecular interaction between Rap1 and Ku are boxed. (**B**) Four *yku80* alleles were tested for NHEJ efficiency (no insert, *yku80Δ* cells complemented with a wild-type or mutant copy of *YKU80* on a *CEN/ARS* vector) and for NHEJ inhibition by Rap1 (one Rap1 site located at 11 bp from the broken end, *rif2Δ sir4Δ yku80Δ* cells complemented with a wild-type or mutant copy of *YKU80* on a *CEN/ARS* vector). See **Fig. 1B** for details of the assay. Mean from independent cell cultures. All four mutants remain NHEJ-proficient and sensitive to Rap1 in this assay. Statistical analysis in **Table S1**.

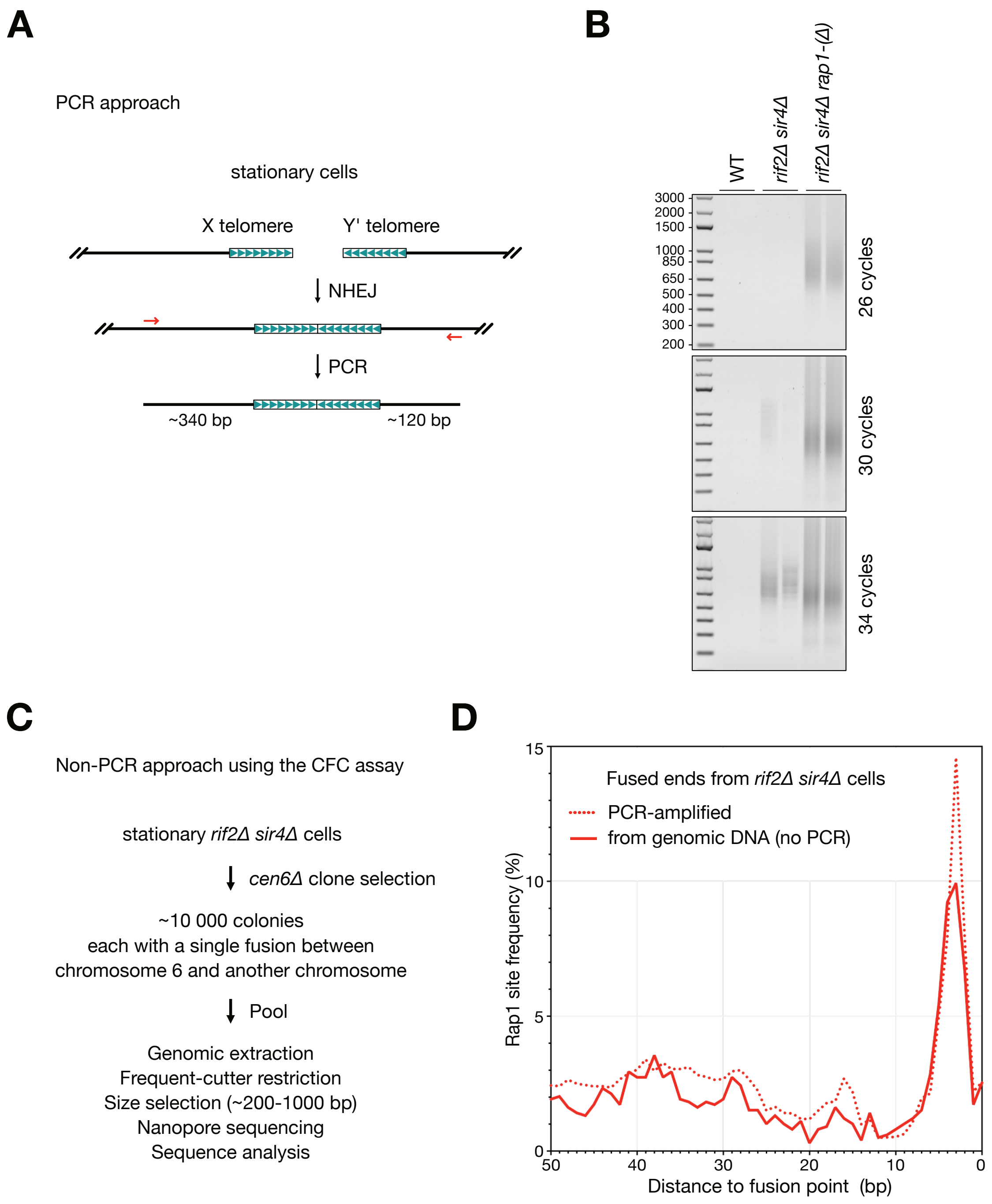


**Supplementary Figure S21. PCR and non-PCR approaches to capture telomere fusions in *S. cerevisiae*.** (**A**) In a PCR approach, telomere fusions were amplified from genomic DNA of stationary cells using a pair of primers hybridizing with X and Y' subtelomeric elements respectively, as described previously (Lescasse et al. 2013). Lengths of the subtelomeric sequences amplified with the fusions are approximately 340 bp from the X elements and 120 bp from the Y' elements. (**B**) Examples of PCR amplifications from *rif2Δ sir4Δ* and *rif2Δ sir4Δ rap1-(Δ)* cells. The number of PCR cycles was adjusted according to the fusion frequencies. PCR products were sequenced by Nanopore sequencing. (**C**) In a non-PCR approach using the Chromosome Fusion Capture assay (CFC) from (Pobiega et al. 2021), we selected a pool of clones carrying a single chromosome fusion from stationary *rif2Δ sir4Δ* cells. The pooled genomic DNA was digested with frequent-cutter restriction enzymes. Telomere sequences, which lack restriction sites, were enriched by size-selection using agarose gel electrophoresis. The selected fragments were sequenced by Nanopore sequencing. (**D**) Rap1 site frequency as a function of the distance to the fusion point in telomere fusions sequenced from genomic DNA without PCR amplification. Dotted line: Rap1 site frequency from PCR-amplified fusions (mean from **Fig. 4B**).

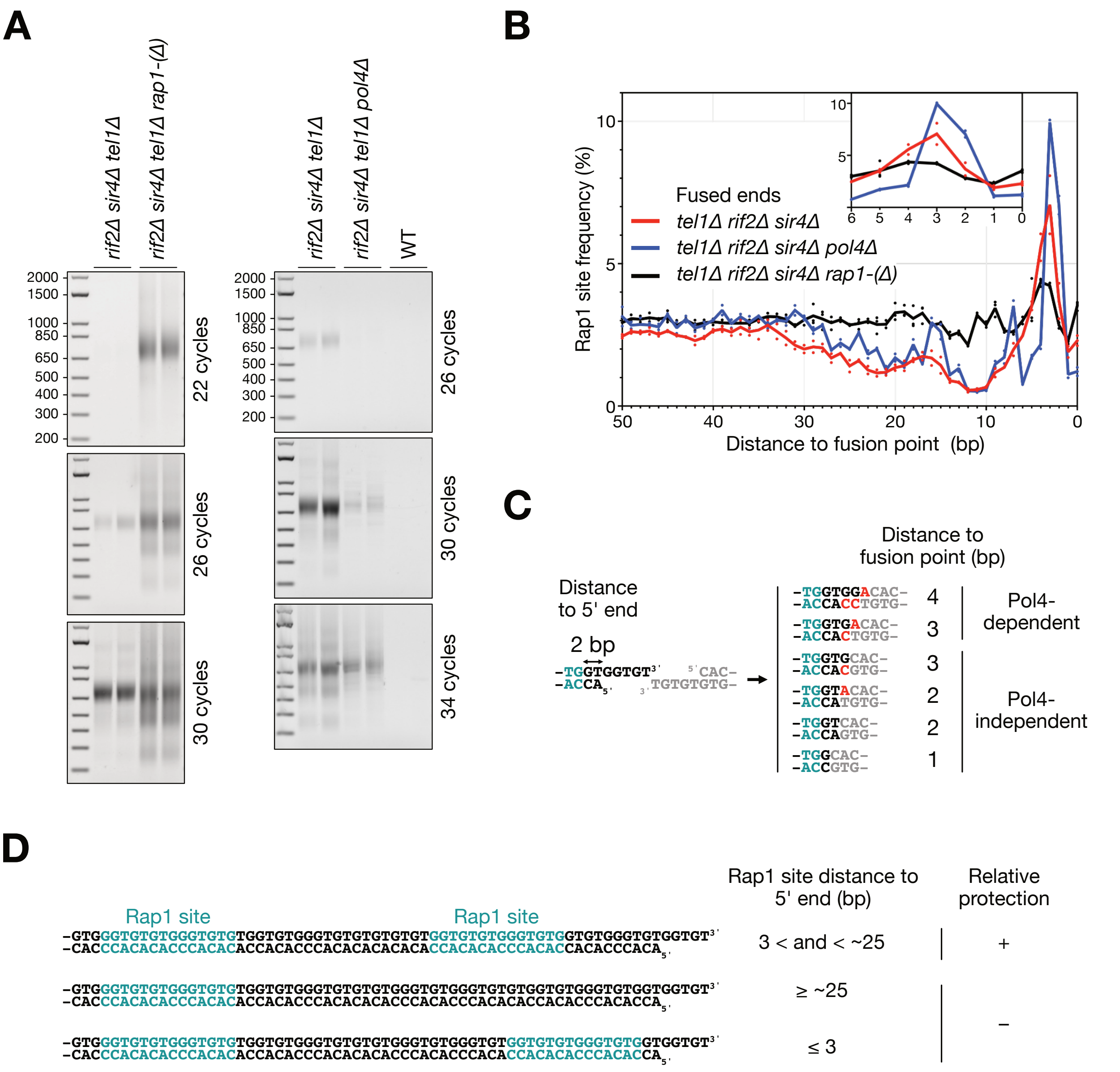


**Supplementary Figure S22. Rap1 site frequency at fused telomere ends in the absence of the NHEJ polymerase Pol4.** (**A**) Examples of PCR amplifications from *rif2Δ sir4Δ tel1Δ*, *rif2Δ sir4Δ tel1Δ rap1-(Δ)* and *rif2Δ sir4Δ tel1Δ pol4Δ* stationary cells. The number of PCR cycles adjusted to the fusion frequencies. PCR products were sequenced by Nanopore sequencing. To compensate for the reduced fusion frequency caused by Pol4 loss (Pardo et al. 2006), we used cells lacking Tel1, whose absence shortens telomeres, thereby further exposing them to NHEJ (Marcand et al. 2008). (**B**) Rap1 site frequency as a function of the distance to the fusion point at fused telomere ends from cells lacking Tel1. Fusion points at position 0. Means from independent cell cultures and sequencing. (**C**) Examples of 3' end trimming and fill-in prior to fusion. The Pol4-dependency was estimated from (Pardo et al. 2006). In red, bases added by Pol4 prior to one strand ligation or by other polymerases after one strand ligation. In teal, the last two base-pairs of a Rap1 site initially 2 bp away from the double-stranded telomeric end. The identity of the enzyme responsible for trimming the overhang remains unknown. (**D**) Examples of telomeres whose position of the end-proximal Rap1 site is either optimum for end protection or too far (≥ 25 bp) or too close (≤ 3 bp) from the telomere double-stranded end. Rap1 sites highlighted in teal.

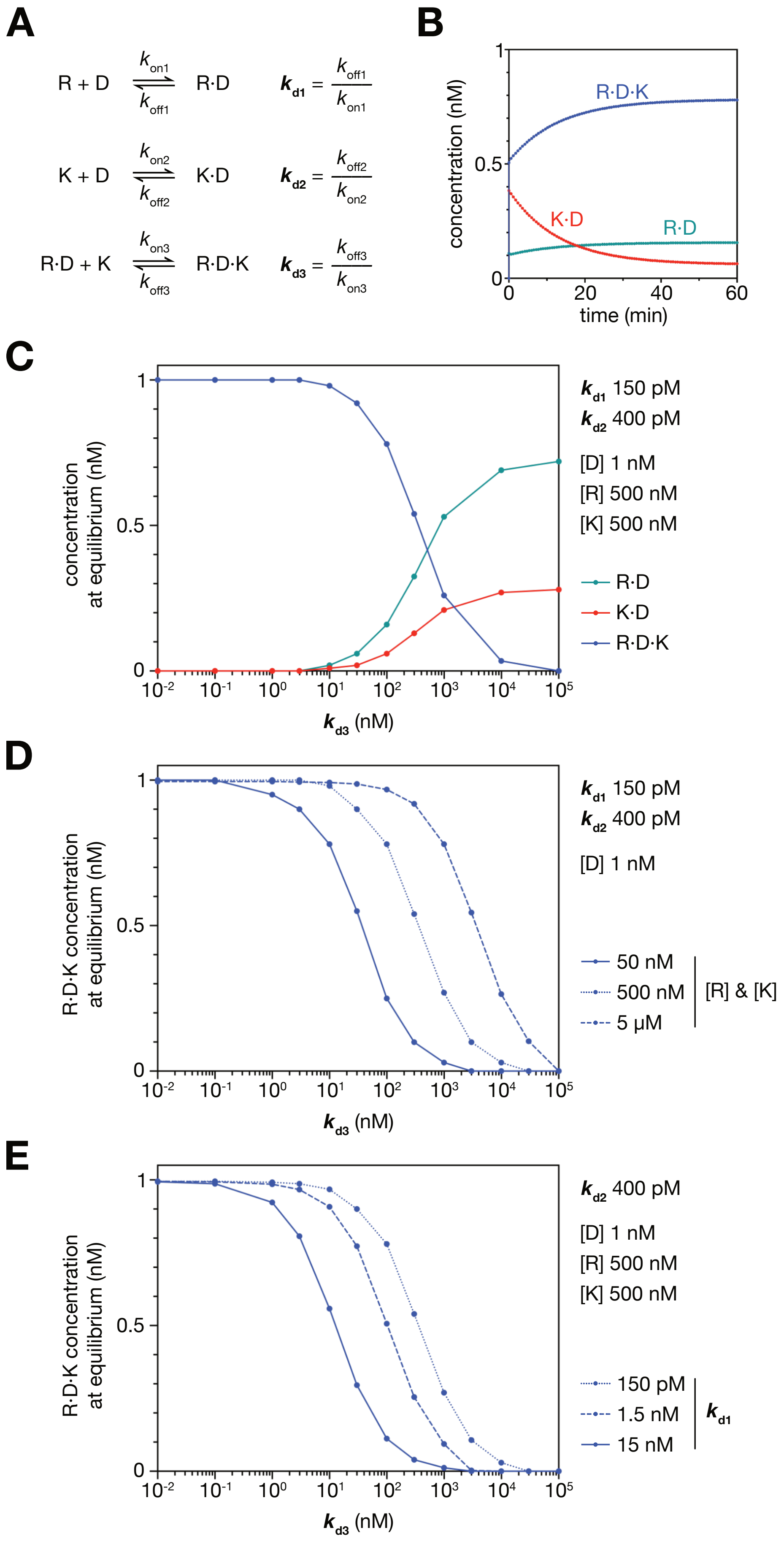
A
R + D ⇌ R·D ($k_{on1}$, $k_{off1}$)
$k_{d1} = \frac{k_{off1}}{k_{on1}}$
K + D ⇌ K·D ($k_{on2}$, $k_{off2}$)
$k_{d2} = \frac{k_{off2}}{k_{on2}}$
R·D + K ⇌ R·D·K ($k_{on3}$, $k_{off3}$)
$k_{d3} = \frac{k_{off3}}{k_{on3}}$
B
concentration (nM)
time (min)
R·D·K
K·D
R·D
C
concentration at equilibrium (nM)
$k_{d3}$ (nM)
$k_{d1}$ 150 pM
$k_{d2}$ 400 pM
[D] 1 nM
[R] 500 nM
[K] 500 nM
R·D
K·D
R·D·K
D
R·D·K concentration at equilibrium (nM)
$k_{d3}$ (nM)
$k_{d1}$ 150 pM
$k_{d2}$ 400 pM
[D] 1 nM
50 nM
500 nM
5 μM
[R] & [K]
E
R·D·K concentration at equilibrium (nM)
$k_{d3}$ (nM)
$k_{d2}$ 400 pM
[D] 1 nM
[R] 500 nM
[K] 500 nM
150 pM
1.5 nM
15 nM
$k_{d1}$

**Supplemental Figure S23**: **Equilibrium binding simulations of the Rap1-DNA-Ku association.** (**A**) Model used to simulate of the equilibrium concentrations. Each equilibrium is characterized by its $k_{off}/k_{on}$ kinetic constants, where $k_{off1}/k_{on1}$ corresponds to the Rap1-DNA binary association (R·D) and $k_{off2}/k_{on2}$ corresponds to the Ku-DNA binary association (K·D). The $k_{d1}$ and $k_{d2}$ values used in these simulations were obtained from the SwitchSENSE experiment shown in **Fig. S3**, except in panel (E). The unknown dissociation constant, $k_{d3}$, for the ternary Rap1-DNA-Ku association (R · D · K) was modelled as the ratio of $k_{off3}/k_{on3}$, with $k_{on3}$ set to $10^7$ $M^{-1}$ $s^{-1}$. The model assumes that steric hindrance revealed by the Cryo-EM structure of the Rap1-DNA-Ku ternary complex prevents direct association/dissociation between Rap1 and Ku-DNA complexes. (**B**) Example of progress curves (from t = 0 to 3600 s) obtained with DynaFit4 according to the model shown in (A) with $[Ku]_0$ = 500 nM, $[Rap1]_0$ = 500 nM, $[DNA]_0$ = 1 nM, $k_{d3}$ = 100 nM ($k_{on3}$ = $10^7$ $M^{-1}$ $s^{-1}$, $k_{off3}$ = 1 $s^{-1}$). (**C**) Evolution of the concentrations of Ku-DNA, Rap1-DNA and Rap1-DNA-Ku complexes as a function of the dissociation constant $k_{d3}$ for the ternary complex dissociating into Ku and Rap1-DNA. The initial concentrations of Ku, Rap1 and DNA are indicated in the left section of the figure. Under these concentrations and given $k_{d1/2}$ values, a $k_{d3}$ of 10 nM or less is sufficient to ensure Rap1-DNA-Ku association at most DNA ends. (**D**) Impact of Ku and Rap1 concentrations (50nM, 500 nM and 5 μM each) on the concentration of the Rap1-DNA-Ku complex. Each curve depicts the Rap1-DNA-Ku concentration as a function of $k_{d3}$ for the different [Ku] and [Rap1] values tested. Lower protein concentrations reduce the likelihood of Rap1-DNA-Ku association. The concentrations of Rap1 and Ku in the nucleus have been estimated to be close to ~1 μM (Ho et al. 2018). (**E**) Effect of Rap1 affinity on the concentration of the Rap1-DNA-Ku complex. Three $k_{d1}$ values (shown on the right side of the graph) were tested. Each curve displays the Rap1-DNA-Ku concentration as a function of $k_{d3}$. Lower Rap1-DNA affinity decreases the likelihood of Rap1-DNA-Ku association.

### REFERENCES


Afonine, P.V., Poon, B.K., Read, R.J., Sobolev, O.V., Terwilliger, T.C., Urzhumtsev, A., and Adams, P.D. (2018). Real-space refinement in *PHENIX* for cryo-EM and crystallography. Acta Crystallogr D Struct Biol *74*, 531–544.

Bordelet, H., et al. (2022). Sir3 heterochromatin protein promotes non-homologous end joining by direct inhibition of Sae2. The EMBO Journal *41*, e108813

Brooks, B.R., Brooks, C.L., Mackerell, A.D., Nilsson, L., Petrella, R.J., Roux, B., Won, Y., Archontis, G., Bartels, C., Boresch, S., et al. (2009). CHARMM: The biomolecular simulation program. J Comput Chem *30*, 1545–1614.

Buchman, A. R., Kimmerly, W. J., Rine, J. and Kornberg, R. D. (1988). Two DNA-Binding Factors Recognize Specific Sequences at Silencers, Upstream Activating Sequences, Autonomously Replicating Sequences, and Telomeres in *Saccharomyces cerevisiae*. Molecular and Cellular Biology *8*, 210–225.

Challal, D. *et al.* (2018) General Regulatory Factors Control the Fidelity of Transcription by Restricting Non-coding and Ectopic Initiation. Molecular Cell *72*, 955-969.e7.

Chan, S.W.L., Chang, J., Prescott, J., and Blackburn, E. H. (2001). Altering telomere structure allows telomerase to act in yeast lacking ATM kinases. Current Biology *11*, 1240–1250.

Chen, H., Xue, J., Churikov, D., Hass, E.P., Shi, S., Lemon, L.D., Luciano, P., Bertuch, A.A., Zappulla, D.C., Géli, V., et al. (2018). Structural Insights into Yeast Telomerase Recruitment to Telomeres. Cell *172*, 331-343.e13.

Eswar, N. (2003). Tools for comparative protein structure modeling and analysis. Nucleic Acids Research *31*, 3375–3380.

Förstemann, K., and Lingner, J. (2001). Molecular Basis for Telomere Repeat Divergence in Budding Yeast. Mol Cell Biol *21*, 7277–7286.

Goddard, T.D., Huang, C.C., Meng, E.C., Pettersen, E.F., Couch, G.S., Morris, J.H., and Ferrin, T.E. (2018). UCSF ChimeraX: Meeting modern challenges in visualization and analysis. Protein Science *27*, 14–25.

Hafner, L., Lezaja, A., Zhang, X., Lemmens, L., Shyian, M., Albert, B., Follonier, C., Nunes, J.M., Lopes, M., Shore, D., et al. (2018). Rif1 Binding and Control of Chromosome-Internal DNA Replication Origins Is Limited by Telomere Sequestration. Cell Reports *23*, 983–992.

Ho, B., Baryshnikova, A., and Brown, G. W. (2018). Unification of Protein Abundance Datasets Yields a Quantitative Saccharomyces cerevisiae Proteome. *Cell Systems 6*, 192-205.e3,

Huang, J., Rauscher, S., Nawrocki, G., Ran, T., Feig, M., De Groot, B.L., Grubmüller, H., and MacKerell, A.D. (2017). CHARMM36m: an improved force field for folded and intrinsically disordered proteins. Nat Methods *14*, 71–73. 10.1038/nmeth.4067.

Humphrey, W., Dalke, A., and Schulten, K. (1996). VMD: Visual molecular dynamics. Journal of Molecular Graphics *14*, 33–38.

Kandiah E., Giraud T., de Maaria Antolinos A., Dobias F., Effantin G., Flot D., Hons M., Schoehn G., Susini J., Svensson O., Leonard G.A. & Mueller-Dieckmann C. (2019). CM01: a facility for cryo-electron microscopy at the European Synchrotron. Acta Cryst. D75, 528 - 535.

Khayat, F. *et al.* (2021) Inhibition of MRN activity by a telomere protein motif. *Nat Commun* 12, 3856.

Kuzmic, P. (1996). Program DYNAFIT for the analysis of enzyme kinetic data: Application to HIV proteinase. Anal. Biochem. 237, 260–273.

Kuzmic, P. (2006). A generalized numerical approach to rapid-equilibrium enzyme kinetics: Application to 17b-HSD. Mol. Cell. Endocrinol. 248, 172–181.

Lescasse, R., Pobiega, S., Callebaut, I., and Marcand, S. (2013). End-joining inhibition at telomeres requires the translocase and polySUMO-dependent ubiquitin ligase Uls1. EMBO J *32*, 805–815.

Liebschner, D., Afonine, P.V., Baker, M.L., Bunkóczi, G., Chen, V.B., Croll, T.I., Hintze, B., Hung, L.-W., Jain, S., McCoy, A.J., et al. (2019). Macromolecular structure determination using X-rays, neutrons and electrons: recent developments in *Phenix*. Acta Crystallogr D Struct Biol *75*, 861–877.

McEachern, M.J., Iyer, S., Boswell Fulton, T., and E.H., Blackburn (2000) Telomere fusions caused by mutating the terminal region of telomeric DNA. PNAS *97*, 11409 –11414

Marcand, S., Pardo, B., Gratias, A., Cahun, S. & Callebaut, I. (2008). Multiple pathways inhibit NHEJ at telomeres. Genes Dev. *22*, 1153–1158.

Matot, B., Le Bihan, Y.-V., Lescasse, R., Pérez, J., Miron, S., David, G., Castaing, B., Weber, P., Raynal, B., Zinn-Justin, S., et al. (2012). The orientation of the C-terminal domain of the Saccharomyces cerevisiae Rap1 protein is determined by its binding to DNA. Nucleic Acids Research *40*, 3197–3207.

Negrini, S., Ribaud, V., Bianchi, A., and Shore, D. (2007). DNA breaks are masked by multiple Rap1 binding in yeast: implications for telomere capping and telomerase regulation. Genes Dev. *21*, 292–302.

Pardo, B., Ma, E. & Marcand, S. (2006). Mismatch Tolerance by DNA Polymerase Pol4 in the Course of Nonhomologous End Joining in *Saccharomyces cerevisiae*. Genetics *172*, 2689–2694

Pettersen, E.F., Goddard, T.D., Huang, C.C., Meng, E.C., Couch, G.S., Croll, T.I., Morris, J.H., and Ferrin, T.E. (2021). UCSF ChimeraX : Structure visualization for researchers, educators, and developers. Protein Science *30*, 70–82.

Pfingsten, J.S., Goodrich, K.J., Taabazuing, C., Ouenzar, F., Chartrand, P., and Cech, T.R. (2012). Mutually Exclusive Binding of Telomerase RNA and DNA by Ku Alters Telomerase Recruitment Model. Cell *148*, 922–932.

Phillips, J.C., Braun, R., Wang, W., Gumbart, J., Tajkhorshid, E., Villa, E., Chipot, C., Skeel, R.D., Kalé, L., and Schulten, K. (2005). Scalable molecular dynamics with NAMD. J Comput Chem *26*, 1781–1802.

Prescott, J.C., and Blackburn, E.H. (2000) Telomerase RNA template mutations reveal sequence specific requirements for the activation and repression of telomerase action at telomeres. Mol. Cell. Biol. 20, 2941-8.

Pobiega, S., Alibert, O., and Marcand, S. (2021). A new assay capturing chromosome fusions shows a protection trade-off at telomeres and NHEJ vulnerability to low-density ionizing radiation. Nucleic Acids Research *49*, 6817–6831.

Šali, A., and Blundell, T.L. (1993). Comparative Protein Modelling by Satisfaction of Spatial Restraints. Journal of Molecular Biology *234*, 779–815.

Roisné-Hamelin, F., Pobiega, S., Jézéquel, K., Miron, S., Dépagne, J., Veaute, X., Busso, D., Du, M.-H.L., Callebaut, I., Charbonnier, J.-B., et al. (2021). Mechanism of MRX inhibition by Rif2 at telomeres. Nat Commun *12*, 2763.

Shen, W., Le, S., Li, Y., and Hu, F. (2016). SeqKit: A Cross-Platform and Ultrafast Toolkit for FASTA/Q File Manipulation. PLoS ONE *11*, e0163962.

Trabuco, L.G., Villa, E., Mitra, K., Frank, J., and Schulten, K. (2008). Flexible Fitting of Atomic Structures into Electron Microscopy Maps Using Molecular Dynamics. Structure *16*, 673–683.

Wriggers, W., Chacon, P. (2001). Modeling Tricks and Fitting Techniques for Multi-Resolution Structures Structure, 9,779–788.

Young, G., Hundt, N., Cole, D., Fineberg, A., Andrecka, J., Tyler, A., Olerinyova, A., Ansari, A., Marklund, E.G., Collier, M.P., et al. (2018). Quantitative mass imaging of single biological macromolecules. Science *360*, 423–427.
